# Structure of mycobacterial NDH-2 bound to a 2-mercapto-quinazolinone inhibitor

**DOI:** 10.1101/2025.01.08.631920

**Authors:** Yingke Liang, Stephanie A. Bueler, Gregory M. Cook, John L. Rubinstein

## Abstract

The mycobacterial type II NADH dehydrogenase (NDH-2) is a promising drug target because of its central role in energy metabolism in *Mycobacterium tuberculosis* and other pathogens, and because it lacks a known mammalian homologue. However, the absence of structural information on how the enzyme binds inhibitors has made optimization of lead compounds challenging. We used electron cryomicroscopy (cryo-EM) to determine the structure of NDH-2 from *Mycobacterium smegmatis*, a fast-growing non-pathogenic model for respiration in *M. tuberculosis*, both alone and in complex with a 2-mercapto-quinazolinone inhibitor. The structure shows that active mycobacterial NDH-2 is dimeric, with the dimerization interface stabilized by an extended C-terminal α helix not found in NDH-2 from other bacterial genera.

The arrangement of monomers in the dimer differs from the arrangement described for other prokaryotic NDH-2 dimers, instead resembling dimers formed by NDH-2 in eukaryotes. Density for the 2-mercapto-quinazolinone in the menaquinone-binding site shows that the inhibitor blocks menaquinone reduction through direct interaction with the flavin adenine dinucleotide cofactor. These results reveal structural elements of NDH-2 that could be used to design specific inhibitors of the mycobacterial enzyme.

## Introduction

The bacterial genus *Mycobacterium* includes several human pathogens, such as *M. leprae, M. abscessus, M. avium,* and most notably *M. tuberculosis*. *M. tuberculosis* is the etiologic agent of the disease tuberculosis (TB), which is the leading cause of death by infectious disease worldwide (World Health Organization, 2023a). Mycobacterial infections require long treatment regimens with multiple antibiotics, which increases the likelihood of developing antibiotic resistance and presents logistic and economic burdens that disproportionately affect developing countries (Dartois and Dick, 2024; World Health Organization, 2023b). Consequently, novel therapeutics and improved treatment regimens are urgently needed.

The success of the F-type adenosine triphosphate (ATP) synthase inhibitor bedaquiline demonstrated the value of oxidative phosphorylation as a target for anti-mycobacterial drug development (Bald et al., 2017; Cox and Laessig, 2014). Oxidative phosphorylation provides cytosolic ATP through the combined activities of the electron transport chain (ETC) and ATP synthase. The main entry point for electrons into the mycobacterial ETC is from the oxidation of reduced nicotinamide adenine dinucleotide (NADH) by NADH:menaquinone oxidoreductases, also known as NADH dehydrogenases. Mycobacteria possess two types of NADH dehydrogenase. The proton-pumping type I NADH dehydrogenase, also known as Complex I, is homologous with mammalian mitochondrial NADH dehydrogenases. The non-proton translocating type II NADH dehydrogenase (NDH-2) does not have a homologue in mammalian mitochondria. While mycobacterial Complex I is expressed primarily when oxygen is abundant and carbon sources in the growth medium are limiting, NDH-2 is expressed under all conditions investigated to date (Berney and Cook, 2010; Cook et al., 2014; Liang et al., 2023). *M. tuberculosis* possesses two paralogues of NDH-2, encoded by the genes *ndh* and *ndhA*, while in *M. smegmatis* has a single gene encoding the protein, *MSMEG_3621* (Vilchèze et al., 2018).

NDH-2 is considered a promising therapeutic target in mycobacteria (Rao et al., 2008; Weinstein et al., 2005; Xu et al., 2023) and other pathogens (Duarte et al., 2021; Lu et al., 2022) because of its important role in the ETC and because there is no human homologue of the protein. CRISPR interference (CRISPRi) gene knockdown of *ndh2* in routine laboratory culture revealed that NDH-2 is bactericidal target in *M. smegmatis* and *M. tuberculosis* (Bosch et al., 2021; McNeil et al., 2022, 2021). Several inhibitors of mycobacterial NDH-2 have already been developed (Harbut et al., 2018; Hong et al., 2017; Korkegian et al., 2018; Nguyen et al., 2018; Shirude et al., 2012), but the utility of these compounds is limited by low specificity, low potency, or poor pharmacokinetic properties. Structure–activity relationship (SAR) studies have been conducted for some inhibitor classes (Dam et al., 2022; Harbut et al., 2018; Murugesan et al., 2018) but the absence of an atomic model of mycobacterial NDH-2 has impeded compound development.

Previous work has determined the structures of many of the enzymes involved in oxidative phosphorylation in mycobacteria: Complex I (Liang et al., 2023), the Rieske protein-containing monomeric succinate dehydrogenase Sdh1 (Zhou et al., 2021), the diheme trimeric succinate dehydrogenase Sdh2 (Gong et al., 2020), the Complex III_2_IV_2_ respiratory supercomplex (Gong et al., 2018; Wiseman et al., 2018), the cytochrome *bd* terminal oxidase (Safarian et al., 2019; Wang et al., 2021), and the ATP synthase (Guo et al., 2021; Zhang et al., 2024). Here we present the structure of mycobacterial NDH-2 determined by electron cryomicroscopy (cryo-EM), providing an important missing piece for a structural model of the mycobacterial ETC. We found that the active form of mycobacterial NDH-2 is a dimer, with the dimer stabilized by a C- terminal α helix that appears unique to mycobacteria. NDH-2 is a peripheral membrane protein, and the cryo-EM map shows a distinct detergent micelle protruding from the membrane-interacting face of the protein, a feature that, to our knowledge, has not been seen previously in cryo-EM maps. Measurement of NADH-driven acidification of native inverted membrane vesicles (IMVs) shows that a 2-mercapto-quinazolinone compound (Murugesan et al., 2018) inhibits NDH-2 at micromolar concentrations. A structure of NDH-2 bound to the 2-mercapto-quinazolinone reveals that this class of inhibitor binds in the putative menaquinone-reducing pocket of the enzyme. Comparison of the structure to structures of NDH-2 from other organisms highlights features in the mycobacterial enzyme that may be important for the rational design of new inhibitors.

## Results

### The active form of mycobacterial NDH-2 is a high molecular weight species

We purified endogenous NDH-2 from a strain of *M. smegmatis* encoding a 3×FLAG sequence in the bacterial genome at the 3′ end of the *ndh* gene (*MSMEG_3621*). NDH-2 is a peripheral membrane protein, and isolation of NDH-2 from other organisms has required detergent solubilization of the membrane before purification (Björklöf et al., 2000; Feng et al., 2012; Heikal et al., 2014; Yang et al., 2017; Yano et al., 2006). Mycobacterial NDH-2 was previously found to be most active in cholates and related detergents, such as deoxycholate and 3-[(3- cholamidopropyl)dimethylammonio]-1-propanesulfonate (CHAPS) (Murugesan et al., 2018; Yano et al., 2006). However, these detergents are usually not optimal for structural studies because of their high critical micelle concentrations (CMCs). Instead, we extracted the protein from the membrane with the low-CMC detergent dodecyl maltoside (DDM), which we used previously with other mycobacterial membrane proteins (Courbon et al., 2023; Guo et al., 2021; Liang et al., 2023; Wiseman et al., 2018; Yanofsky et al., 2021). *M. smegmatis* NDH-2 was purified by anti-FLAG affinity chromatography followed by size exclusion chromatography (SEC). The SEC chromatogram (**Fig. S1A**) showed a large peak that eluted at low volume, which we subsequently found contained ribosomes, as well as two poorly-resolved peaks that eluted at higher volume. Sodium dodecyl sulfate polyacrylamide gel electrophoresis (SDS-PAGE) revealed that both partially resolved peaks result from an ∼50 kDa polypeptide (**Fig. S1B**, *black arrow*), corresponding to the expected 49 kDa mass of the *M. smegmatis* NDH-2. Assay of SEC fractions using the soluble electron acceptor decylubiquinone found NADH:quinone oxidoreductase activity only for the partially-resolved peak that eluted at lower volume, corresponding to a higher molecular weight species (**Fig. S1C**). These results suggested that mycobacterial NDH-2 could exist both as a dimer and as a monomer, but with only the dimer being active. In *S. cerevisiae*, NDH-2, dimerization was found to be critical for catalysis and membrane anchoring (Feng et al., 2012). However, the reason why dimerization is necessary for function could not be inferred from the structure. Similarly, all previous structures of NDH-2 from both prokaryotes and eukaryotes were dimeric (Feng et al., 2012; Heikal et al., 2014; Sena et al., 2015; Yang et al., 2017).

### Mycobacterial NDH-2 functions as a dimer that is stabilized by a C-terminal α helix

By exchanging NDH-2 into different detergents and imaging these specimens by cryo-EM, we found that samples in the detergent lauryl maltose neopentyl glycol (LMNG) gave images with a higher density of protein particles than the same concentration of protein in DDM. Therefore, SEC fractions with NADH:quinone oxidoreductase activity were exchanged into LMNG and subjected to structural analysis by cryo-EM (**Fig. S2A**). *Ab initio* three-dimensional (3D) reconstruction followed by iterative map refinement with C2 symmetry allowed calculation of a map at a nominal overall 3.0 Å resolution (**Fig. 1A**, *upper*, **Fig. S3**, **Table S1**). The map showed a dimer of NDH-2 at sufficient resolution to build an atomic model for 98 % of the residues in the complex (**Fig. 1A**, *lower*, **Table S1**). The model includes a single flavin adenine dinucleotide (FAD) molecule in each protomer as the only redox active cofactor (**Fig. 1B**). Interestingly, class average images from two-dimensional (2D) classification show a clear detergent micelle (**Fig 1C**, *white triangles*, **Fig. S3A**, *lower*). This micelle is also visible in the cryo-EM map at low threshold (**Fig. 1D**, *gray surface*), marking the hydrophobic face of the protein and defining its orientation with respect to the membrane. Only a few cryo-EM structures of peripheral membrane proteins have been published previously. Most of these structures are intermediate-resolution maps of membrane remodelling complexes that transition from a soluble state to a membrane-bound state following a change in environment (Dodonova et al., 2017; Kovtun et al., 2018, 2020; Pang et al., 2019). Recently, a high-resolution cryo-EM structure of the membrane-interacting protein AP2 was determined by mixing pre-formed lipid nanodiscs with AP2 (Cannon et al., 2023). In contrast to earlier structures of peripheral membrane proteins, the NDH- 2 structure shows how detergents interact with a peripheral membrane protein, and that the detergent-solubilized protein can be subjected directly to structural studies by cryo-EM.

**Figure 1:**
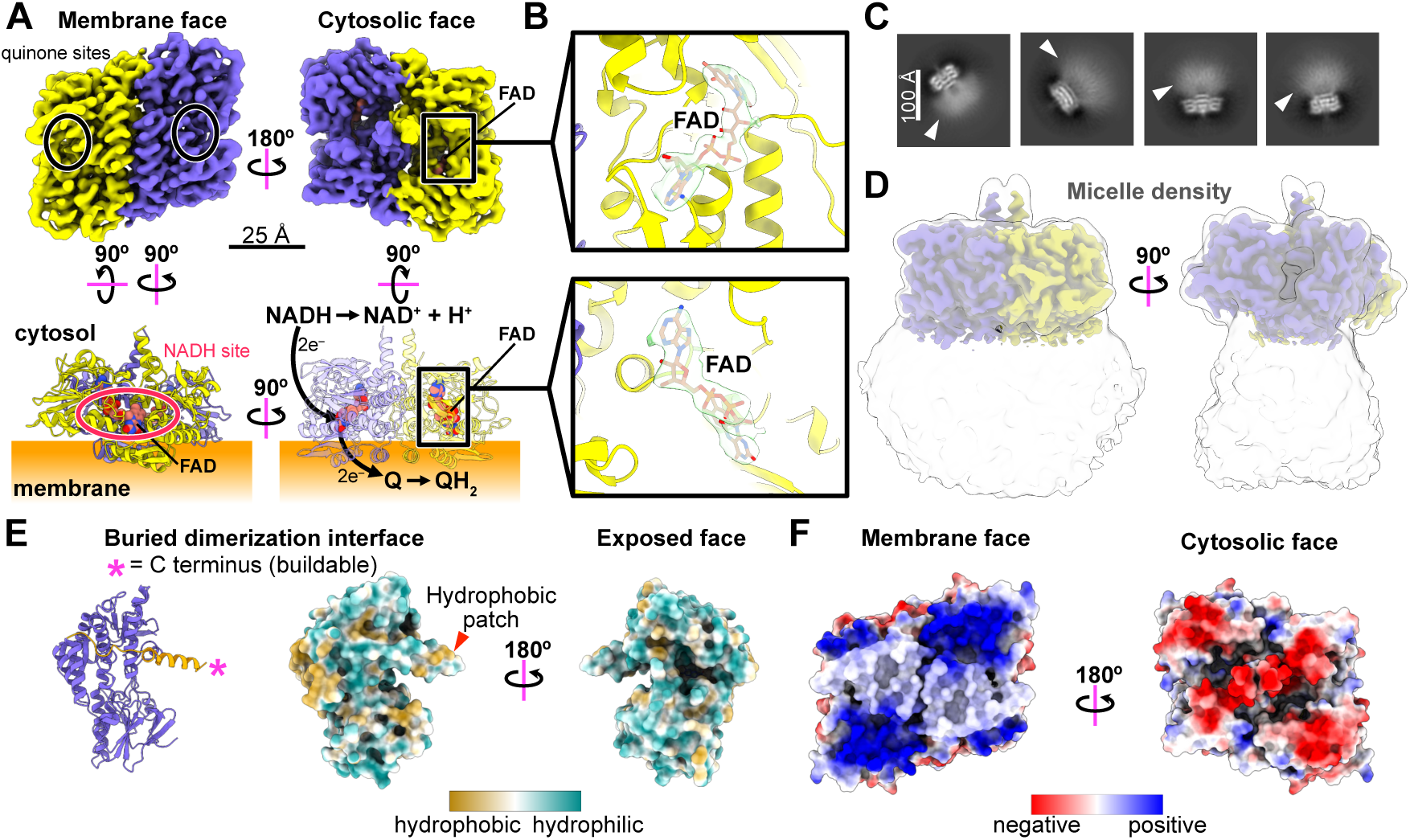
Structure of *M. smegmatis* NDH-2. **A)** Cryo-EM map (*top*) and atomic model (*bottom*) of NDH-2, with the membrane indicated (*orange rectangle*). The positions of the quinone-binding sites (*top left, black ovals*), an NADH-binding site (*bottom left, red oval*), and a flavin adenine dinucleotide (FAD) cofactor (*right*, *black rectangles*) are indicated. The electron transfer path is shown in the bottom right panel. **B)** Close-up view of FAD cofactor in the cryo-EM map. **C)** Selected 2D class average images with detergent micelles indicated by white triangles. **D)** Cryo-EM map showing the detergent micelle (*gray surface*) extending from the membrane face of NDH-2. **E)** *Left*: View of NDH-2 monomer highlighting the C-terminal helical extension (*orange helix*). The model was built for residues 2 to 447 out of a total of 457 residues. Residue 447 is indicated with an asterisk (*). *Centre*: Surface hydrophobicity for the dimerization interface of the NDH-2 monomer, highlighting the hydrophobic patch on the C- terminal α helix (*red triangle*). *Right*: Surface hydrophobicity for the face of the same monomer that interacts with the cytosol. **F)** Electrostatic surface of the NDH-2 dimer.

The two protomers in the dimer have a large (∼1634 Å^2^) interaction interface (Krissinel and Henrick, 2007). Like NDH-2 from other organisms (Feng et al., 2012; Heikal et al., 2014; Sena et al., 2015; Yang et al., 2017), the mycobacterial NDH-2 protomer is an α/β protein composed of an N-terminal active domain (residues 11–349) joined to a C-terminal domain (residues 366– 457) by a linker. The active domain consists of two Rossmann-like folds that form two nucleotide binding sites: one for the substrate NADH in a solvent-accessible pocket (**Fig. 1A**, *bottom left, red oval*) and the other for the FAD cofactor near the centre of the protein (**Fig 1A**, *bottom right*, *black square*). The menaquinone binding site is located where NDH-2 interacts with the membrane (**Fig. 1A**, *top left*, *black ovals*). The C-terminal sequence of mitochondrial NDH-2 forms a domain that has been suggested to be important for oligomerization (Feng et al., 2012). In the mycobacterial enzyme, the C-terminal sequence of each protomer (residues 430– 457) forms an α helix at the center of the dimer that spans from membrane to dimer’s cytoplasmic face (**Fig. 1E**, *left, orange α helix*). This α helix contains a hydrophobic patch that helps stabilize the dimer as part of the dimer interface (**Fig. 1E**, *middle, red triangle*). The structure of mycobacterial NDH-2 is consistent with a proposed mechanism where electrons are transferred from NADH to menaquinone via the FAD cofactor in a two-step process (Melo et al., 2004) with the binding of NADH and quinone independent of each other (Blaza et al., 2017) (**Fig. 1A**, *bottom right*).

Mycobacteria have a unique cytosolic membrane, with the inner leaflet comprised almost exclusively of acylphosphatidylinositol dimannoside and diacylphosphatidylinositol dimannoside (Bansal-Mutalik and Nikaido, 2014). These lipids give the inner leaflet a negative surface charge that is complemented by large patches of positive charge on the membrane face of NDH-2 (**Fig. 1F**, *left*). The extensive positive charge on the membrane face appears to be unique to mycobacterial NDH-2 (**Fig. S4A**). In contrast, the cytosolic face of mycobacterial NDH-2 is negatively charged (**Fig. 1F**, *right*).

### The NDH-2 structure is conserved among mycobacteria

Prediction of the structures of the two *M. tuberculosis* NDH-2 paralogues with AlphaFold2 (Jumper et al., 2021; Varadi et al., 2022) suggests that they both closely resemble the structure of the *M. smegmatis* enzyme determined here (**Fig. 2A**). Mapping sequence conservation for homologs from *M. tuberculosis*, *M. leprae*, *M. abscessus*, and *M. avium* onto the structure of the *M. smegmatis* enzyme shows several regions that are highly conserved. Most notably, these regions include the NADH- and menaquinone-binding sites (**Fig. 2B**). This observation suggests that the structure presented here can be used as a model for rational design of inhibitors that block NDH-2 activity in several pathogenic mycobacteria.

**Figure 2:**
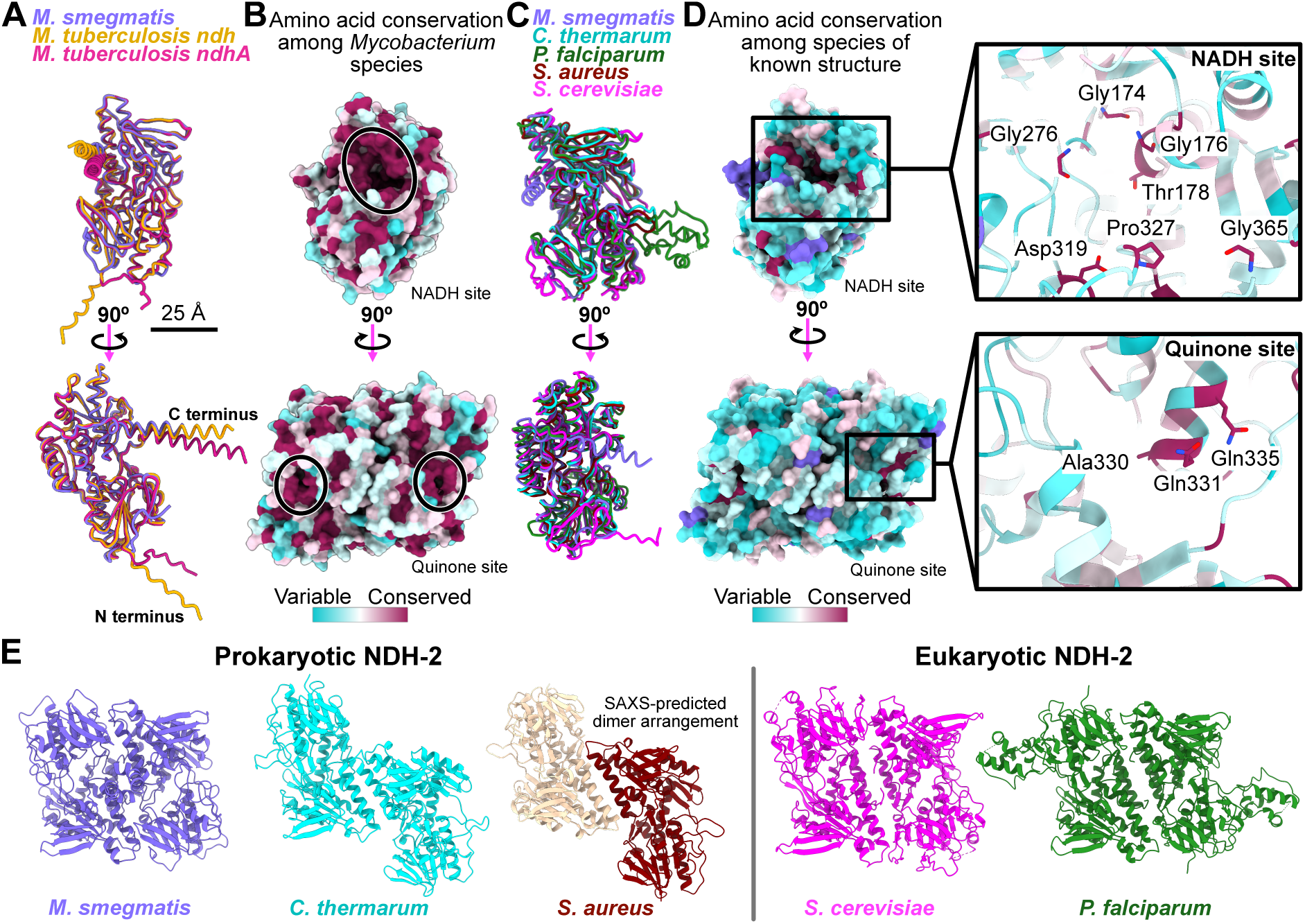
Comparison of *M. smegmatis* NDH-2 with NDH-2 from other species. **A)** Aligned models of *M. smegmatis* NDH-2 monomer with AlphaFold2-predicted models of *M. tuberculosis* ndh and ndhA. **B)** Amino acid conservation between *M. smegmatis* NDH-2 and NDH-2 from *M. tuberculosis*, *M. leprae*, *M. avium*, and *M. abscessus*. Regions of sequence conservation at the quinone and NADH-binding sites are indicated (*ovals*). **C)** Alignment of the *M. smegmatis* NDH-2 monomer structure with experimental structures of NDH-2 from *C. thermarum* (PDB: 6BDO), *P. falciparum* (PDB: 5JWB), *S. aureus* (PDB: 5NA1), and *S. cerevisiae* (PDB: 4G74). **D)** *Left*: Amino acid conservation between *M. smegmatis* NDH-2 and other experimental NDH-2 structures. *Right*: Conserved NADH-binding (*top*) and quinone-binding (*bottom*) residues. **E)** Comparison of prokaryotic (*left*) and eukaryotic (*right*) NDH-2 dimers.

*The mycobacterial NDH-2 dimer resembles eukaryotic, but not other prokaryotic, NDH-2 dimers* Structures of NDH-2 have been determined previously from the eukaryotes *Saccharomyces cerevisiae* (Feng et al., 2012) and *Plasmodium falciparum* (Yang et al., 2017), and the prokaryotes *Caldalkalibacillus thermarum* (Heikal et al., 2014) and *Staphylococcus aureus* (Sena et al., 2015). Comparison of the structure of the mycobacterial NDH-2 monomer with the previously determined structures shows a conserved fold at the core of the protein (**Fig. 2C**).

Similarly, several residues at the NADH-binding site are conserved in all the structures (**Fig. 2D**, *top*). However, unlike NDH-2 from mycobacteria, few residues are conserved at the quinone-binding site (**Fig. 2D**, *bottom*). The most striking difference between known NDH-2 structures is the arrangement of protomers in the dimer. The mycobacterial NDH-2 dimer appears more like eukaryotic NDH-2 dimers than prokaryotic NDH-2 dimers (**Fig. 2E**). This discrepancy may be due to differences in the second of the two membrane-anchoring α helices in the enzyme, which is longer in *M. smegmatis* and the eukaryotic structures than in the prokaryotic structures (**Fig S4B**, *black box*). This membrane-anchoring α helix forms a portion of the dimerization interface in the mycobacterial and eukaryotic NDH-2s, while in *C. thermarum* and *S. aureus* NDH-2 the corresponding, but shorter, α helices may not be able to dimerize in the same way. However, we cannot exclude the possibility that the dimerization interfaces observed in the X-ray crystallography structures of *C. thermarum* and *S. aureus* NDH-2 are the result of crystal packing. The longer α helices also result in a larger hydrophobic domain for the mycobacterial and eukaryotic enzymes (**Fig. S4C**). Interestingly, the C-terminal α helix of mycobacterial NDH- 2, which also participates in dimerization, is not found in other experimentally determined NDH- 2 structures (**Fig. S4D**, *green triangles*). The computer program Foldseek (van Kempen et al., 2024) suggests that NDH-2 from *Trypanosoma cruzi* and *T. brucei* are the most closely related proteins with a similar C-terminal α helix.

### A 2-mercapto-quinazolinone inhibits NDH-2 activity in native membranes

To characterize inhibitors that bind NDH-2 in its native lipid bilayer, we used NADH to drive acidification of IMVs prepared from *M. smegmatis* cytosolic membranes (Koul et al., 2007) and quantified acidification by measuring fluorescence recovery after adding an ionophore (Harden et al., 2024). In this assay, IMVs are incubated with the fluorophore 9-amino-6-chloro-2- methoxyacridine (ACMA) and NADH is added as an electron donor. The NADH:menaquinone oxidoreductase activity of NDH-2 provides menaquinol to the ETC. This menaquinol is oxidized by the Complex III_2_IV_2_ supercomplex and cytochrome *bd*, which translocate protons, acidifying the IMV lumen and quenching ACMA fluorescence. NDH-2 inhibitors are expected to block NADH-driven IMV acidification, which can be measured from the fluorescence recovery after adding the H^+^/K^+^ antiporter nigericin. Specific inhibition of NADH:menaquinone oxidoreductase activity can be distinguished from inhibition of other ETC complexes or non-specific uncoupling of the proton motive force with two control experiments. In the first control experiment, succinate is used to drive IMV acidification by providing electrons to the succinate:menaquinone oxidoreductases (Pecsi et al., 2014). For the second control experiment, ATP is used to drive acidification of IMVs from an *M. smegmatis* strain where the ATP synthase has been modified to allow it to function as a proton pump (Guo et al., 2021; Harden et al., 2024). Inhibition of both NADH- and succinate-driven IMV acidification, but not ATP-driven IMV acidification, would suggest that a compound targets menaquinol oxidation. Compounds that block NADH-, succinate-, and ATP-driven acidification are likely non-specific inhibitors or ionophores (Nakatani et al., 2020). In contrast, inhibition of only NADH-driven acidification but not succinate- or ATP-driven acidification suggests that a compound is a specific inhibitor of NADH:menaquinone oxidoreductase activity.

Assay of NDH-2 activity with two previously reported inhibitors, quinolinyl pyrimidine Compounds **13A** and **13C** (Shirude et al., 2012) (**Fig. 3A** and **3B**, *top*), indicated inhibition of NADH- (**Fig. 3A** and **3B**, *blue bars*, **Fig. S5**), succinate- (**Fig. 3A** and **3B**, *bottom, magenta bars,* **Fig. S5**), and ATP-driven acidification of the IMVs (**Fig. 3A** and **3B**, *bottom, yellow bars,* **Fig. S5**). The plots show the mean ± std for n = 3 technical replicates, and an independent set of assays is shown in **Fig. S6**. These data suggest that Compounds **13A** and **13C** inhibit IMV acidification by non-specific uncoupling of the proton motive force. While this finding does not preclude inhibition of NDH-2 by **13A** and **13C**, it indicates that if inhibition occurs, it occurs above the concentration where the compounds induce uncoupling. Non-specific uncoupling of the proton-motive force, which could also affect mitochondria, may explain why many quinolinyl pyrimidines are cytotoxic (Lu et al., 2022).

**Figure 3:**
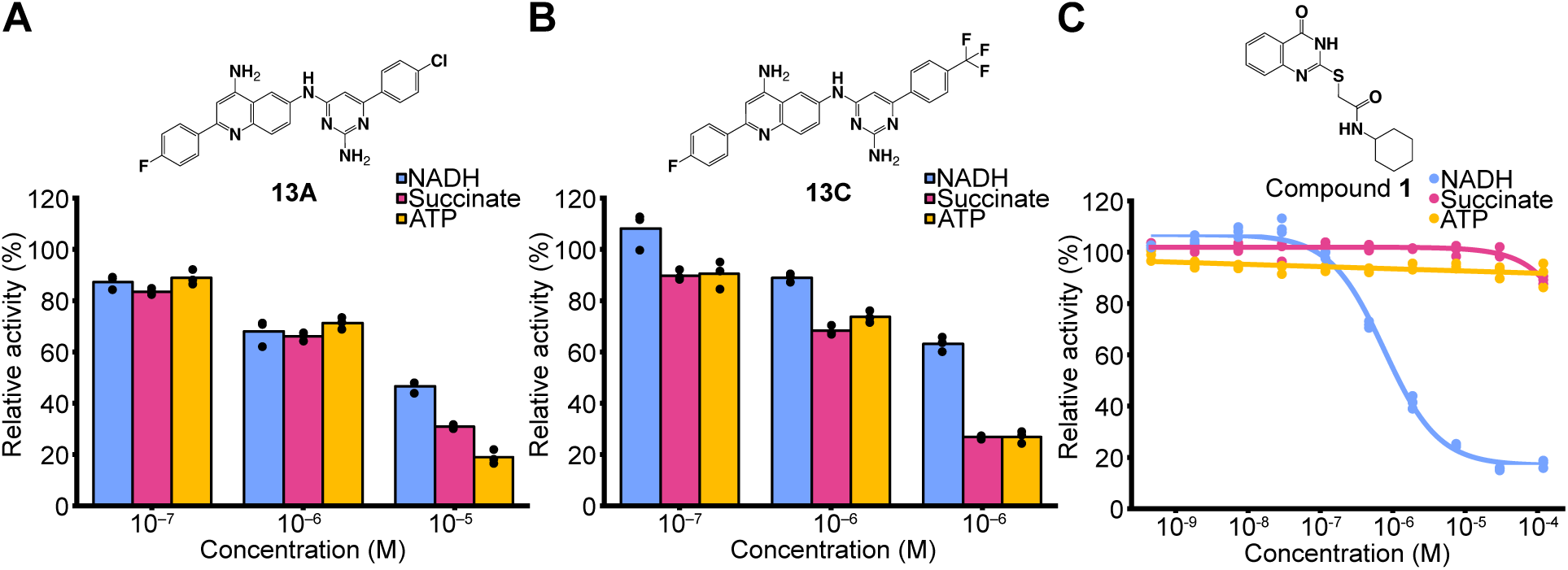
Characterization of mycobacterial NDH-2 inhibitors. **A)** NADH- (*blue*), succinate- (*magenta*), and ATP-driven (*yellow*) IMV acidification at different concentration of quinolinyl pyrimidine Compound **13A** (*top*). **B)** NADH- (*blue*), succinate- (*magenta*), and ATP-driven (*yellow*) IMV acidification at different concentration of quinolinyl pyrimidine Compound **13C** (*top*). **C)** Dose-response curve for NADH- (*blue*), succinate- (*magenta*), and ATP-driven (*yellow*) IMV acidification at different concentrations of 2-mercapto-quinazolinone Compound **1** (*top)*.

In addition to the quinolinyl pyrimidines, 2-mercapto-quinazolinone compounds were identified as NDH-2 inhibitors in a screen against *M. tuberculosis* in liquid culture (Murugesan et al., 2018). We tested inhibition of NADH-, succinate-, and ATP-driven IMV acidification by the lead 2-mercapto-quinazolinone, named Compound **1** (2-mercapto-quinazolinone N-cyclohexyl- 2-[(4-oxo-1H-quinazolin-2-yl)sulfanyl]acetamide), which was one of the most potent compounds in the series (**Fig. 3C**, *top*, **Fig****. S6**). Compound **1** inhibited NADH-driven IMV acidification with a micromolar IC_50_ (**Fig. 3C**, *blue curve*, **Fig. S5**, **Fig. S6**). The plot shows the mean ± std for n = 3 technical replicates, and an independent assay is shown in **Fig. S6**. This micromolar IC_50_ is somewhat higher than the IC_50_ of 23 ± 8 nM determined previously using an NADH:Q2 oxidoreductase assay with the purified enzyme (Murugesan et al., 2018), which may be due to differences in the assays. Partial inhibition of succinate- and ATP-driven IMV acidification by Compound **1** was detected only above 10 µM, indicating that Compound **1** is a specific inhibitor of NADH:menaquinone oxidoreductase with little non-specific or off target uncoupling activity (**Fig. 3C**, *magenta and yellow curves*, **Fig. S5**, **Fig. S6**).

### *2-mercapto-quinazolinones* inhibit NDH-2 by blocking the quinone-binding site

Compound **1** consists of a 2-mercapto-quinazolinone headgroup (**Fig. 4A**, *blue*) and a tail composed of an amide moiety (**Fig. 4A**, *magenta*) and a cyclohexyl moiety (**Fig. 4A**, *yellow*). To determine how **1** inhibits NDH-2, we incubated LMNG-solubilized NDH-2 with 25 μM Compound **1**, prepared and imaged cryo-EM specimens, and calculated a 3D map of the protein-inhibitor complex (**Fig. S2B**). Three-dimensional map refinement with C2 symmetry enforced led to a map with a nominal overall resolution of 3.0 Å (**Fig. 4B**, *upper*, **Fig. S7**), which was sufficient to build an atomic model for 97% of residues in the complex (**Fig. 4B**, *lower*, **Table S1**). This 3D map revealed density corresponding to **1** in the menaquinone-binding site of each NDH-2 monomer (**Fig. 4B**, *upper*, **Fig. 4C**, *green surface and model*).

**Figure 4:**
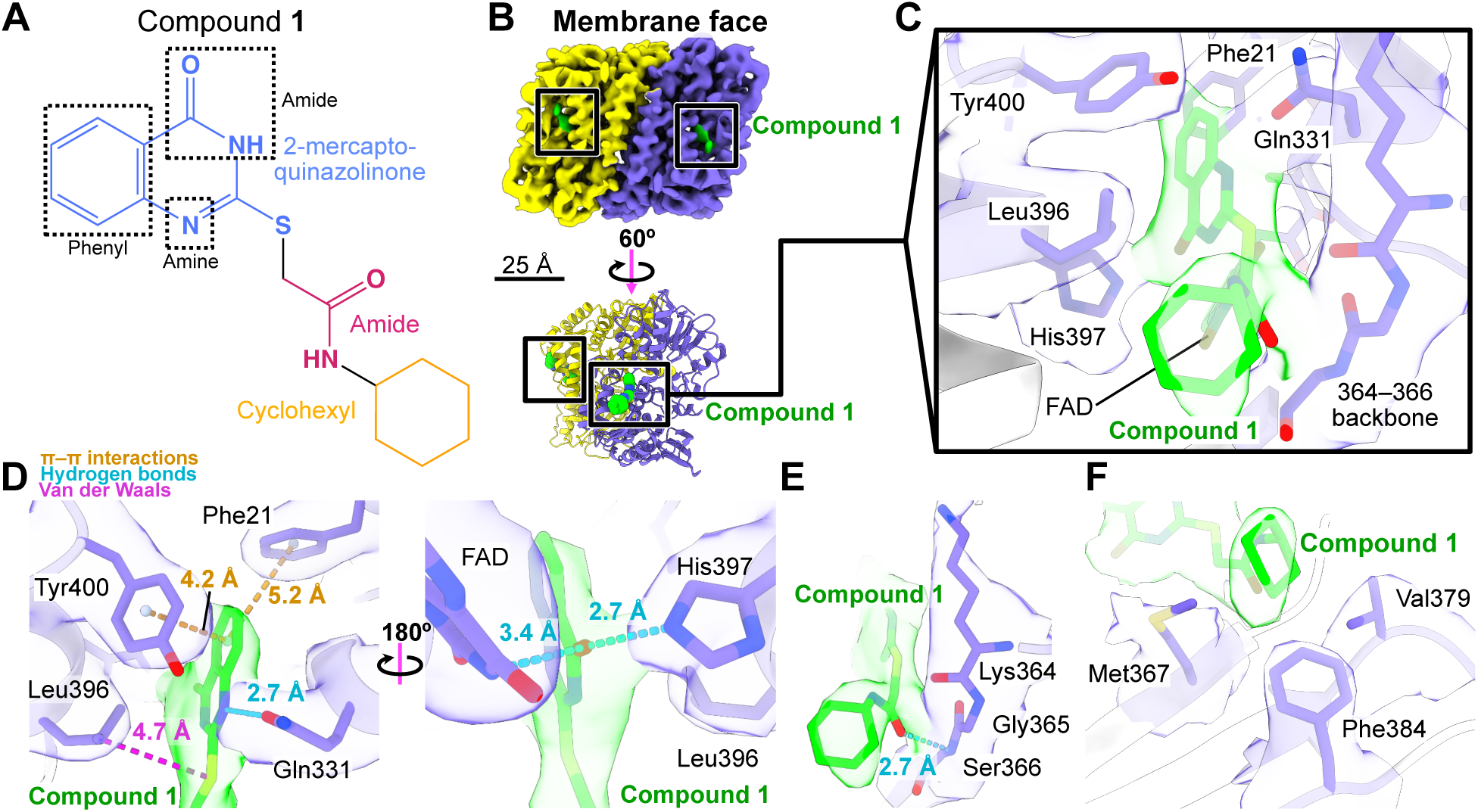
Structure of a 2-mercapto-quinazolinone-bound *M. smegmatis* NDH-2. **A)** Structure of 2-mercapto-quinazolinone Compound **1**. **B)** Cryo-EM map (*top*) and atomic model (*bottom*) for the protein-inhibitor complex. Compound **1** is shown in green. **C)** The Compound **1** binding pocket. **D)** Close-up view of interactions between NDH-2 and the 2-mercapto-quinazolinone moiety of **1**. Intermolecular distances for apparent π–π interactions (*orange dashed lines*), hydrogen bonds (*cyan dashed lines*), and hydrophobic interactions (*magenta dashed line*) are indicated. **E)** Close-up view of interactions between NDH-2 and the tail amide of Compound **1**. Interatomic distance of an apparent hydrogen bond (*cyan dashed line*) is indicated. **F)** Interactions of NDH-2 with the cyclohexyl moiety of Compound **1**.

Most of the interactions between NDH-2 and Compound **1** appear to involve the 2-mercapto-quinazolinone headgroup of the inhibitor (**Fig. 4C**). Tyr400 and Phe21 engage in apparent π–π interactions with the phenyl moiety of the quinazolinone ring, with centre-to-centre distances between the residues and the ring within the range typically observed in protein-ligand interactions (Brylinski, 2017; Burley and Petsko, 1985) (**Fig. 4D**, *left, orange dashed lines*).

Possible hydrogen bonds (**Fig. 4D**, *cyan dashed lines*) between NDH-2 and **1** appear to be mediated primarily by Gln331 and His397, which interact with the amine nitrogen and amide oxygen of the quinazolinone, respectively, from opposite sides of the inhibitor binding pocket. The FAD cofactor from NDH-2 may also form a hydrogen bond with the amide oxygen from **1**, although the distance between these two groups is larger. Leu396 appears to engage in a hydrophobic Van der Waals interaction with the sulfur of the 2-mercapto-quinazolinone moiety (Lindorff-Larsen et al., 2012; Onofrio et al., 2014) (**Fig. 4D**, *magenta dashed line*). The large number of possible interactions between NDH-2 and the 2-mercapto-quinazolinone headgroup highlight the potential of related compounds as inhibitors.

The tail portion of Compound **1** also appears to interact with NDH-2. The amide oxygen from the tail forms a possible hydrogen bond with the backbone nitrogen of Ser366 (**Fig. 4E**, *cyan dashed line*). Furthermore, the orientation of Lys364 puts the backbone oxygen from the residue within hydrogen bonding distance of the amide nitrogen from the tail of **1** (**Fig. 4E**). Finally, the cyclohexyl moiety of **1** form apparent Van der Waals interactions with several hydrophobic residues, notably Met367, Phe384, and Val379, which are all pointed toward **1** and have intermolecular distances within 4 Å, forming a pocket (**Fig. 4F**).

## Discussion

The arrangement of protomers in the mycobacterial NDH-2 structure presented here highlights differences between mycobacterial NDH-2 and NDH-2 from other bacteria. Furthermore, comparison of amino acid conservation between species with known NDH-2 structures reveals that while NADH-binding sites are conserved, quinone-binding sites are highly divergent.

Therefore, inhibitors that bind in the NADH site may be less species-specific than inhibitors that bind the quinone site. Previously, a structure was determined for the inhibitor 2-heptyl-4- hydroxyquinoline-N-oxide (HQNO) bound to NDH-2 from *C. thermarum* (Petri et al., 2018).

HQNO was also found in the quinone-binding site of NDH-2, but was not as deep in the site as Compound **1** (**Fig. S8A**). As a result, HQNO is positioned closer to the FAD cofactor than Compound **1**. Only residues Gln317, Ile379, and FAD in *C. thermarum* NDH-2 were found to play a role in HQNO binding, and interacted only with the 4-hydroxyquinoline N-oxide headgroup (Petri et al., 2018) (**Fig. S8B**). The relatively few contacts between HQNO and *C. thermarum* NDH-2 may explain why HQNO inhibits quinol-binding enzyme non-specifically (Van Ark and Berden, 1977), while Compound **1** appears specific to mycobacterial NDH-2 (Murugesan et al., 2018).

The Compound **1**-bound NDH-2 structure helps rationalize a SAR study that found substantially higher MIC_99_s for several derivatives of Compound **1** (Murugesan et al., 2018). First, substitution of the cyclohexyl moiety with smaller groups could decrease surfaces available for interactions with the hydrophobic pocket formed by Met367, Phe384, and Val379. Addition to positions 2 or 3 of the cyclohexyl moiety, or substitution of cyclohexane with a less flexible group could create steric clashes with the protein. All modifications of the amide in the tail of **1** that were tested involved adding a methyl group, which could disrupt hydrogen bonding with the backbone of residues 364 to 366, as well as cause steric clashes. Substitution of the sulfur in position 2 of the 2-mercapto-quinazolinone moiety with the less polarizable carbon, nitrogen, or oxygen increased the MIC_99_, possibly by interfering with the compounds’ abilities to engage in Van der Waals interaction with Val379 in NDH-2. The quinazolinone moiety of **1** appears to engage in several π–π interactions and hydrogen bonds with NDH-2, explaining why substitution of this group generally led to increased MIC_99_ and reduced potency. However, fluorination of the quinazolinone phenyl moiety appeared to preserve the MIC_99_ while improving compound stability. Therefore, modifying the quinazolinone with aromatic directing groups may be of interest.

The 2-mercapto-quinazolinone compounds have been proposed to be allosteric inhibitors that affect binding of both NADH and menaquinone (Murugesan et al., 2018). However, the results presented here suggest that Compound **1** binds the menaquinone-reducing site of mycobacterial NDH-2, and its binding mode can explain the SAR analysis performed previously. Compound **1** and other 2-mercapto-quinazolinones show promise as specific, non-cytotoxic, and chemically versatile inhibitors of both Ndh and NdhA in *M. tuberculosis* (Harbut et al., 2018; Murugesan et al., 2018). Despite this promise, the utility of these compounds is currently limited by their instability in cells, which is believed to be mediated by glutathione-dependent cleavage at the sulfur atom in the 2-mercapto-quinazolinone moiety (Murugesan et al., 2018). The structure presented here may guide modification of the compounds to improve their stability in cells without compromising binding to their target. Another recently identified class of mycobacterial NDH-2 inhibitors, tricyclic spirolactams, appeared to display allosteric inhibition of NDH-2 from *M. marinum* that was similar to Compound **1**. This inhibition was inactivated by mutating menaquinone binding site residues Tyr403 and Gln334 (Dam et al., 2022), which correspond to Tyr400 and Gln 331 in the *M. smegmatis* enzyme. This observation may indicate that tricyclic spirolactams also bind to the menaquinone site.

Genetic analysis has shown that inactivation of either *ndh* or *<u>ndhA</u>* is not lethal in *M. tuberculosis* (Beites et al., 2019; Vilchèze et al., 2018). However, simultaneous inactivation of both *ndh* and *<u>ndhA</u>* is lethal when *M. tuberculosis* is grown in medium that includes fatty acids, but not in fatty acid-free conditions (Beites et al., 2019). Even in the absence of fatty acids, the double mutant is sensitive to the Complex I inhibitor rotenone, which kills the bacteria by depleting the NAD^+^ available as an electron acceptor (Beites et al., 2019; Xu et al., 2023). Mutations in NDH-2 have previously been shown to confer isoniazid resistance in *M. tuberculosis*, *M. smegmatis*, and *M. bovis* BCG (Lee et al., 2001; Miesel et al., 1998; Vilchèze et al., 2005). However, recent work found that genetic inactivation of *M. tuberculosis ndh* did not confer resistance to isoniazid or other inhibitors of cell wall synthesis (Pandey et al., 2023). In fact, deletion of the gene for NDH-2 was found to sensitize *M. tuberculosis* to killing by bedaquiline in laboratory culture, suggesting that NDH-2 inhibitors could enhance the potency of existing therapeutics (Beites et al., 2019). NDH-2 is the only NADH dehydrogenase in *M. leprae*, the causative agent of leprosy, making NDH-2 inhibitors particularly interesting for the treatment of this disease. Leprosy is a neglected tropical disease (Chen et al., 2022; White and Franco-Paredes, 2015) with ∼200,000 new cases detected each year, and with emerging drug resistance (World Health Organization, 2023b). Therefore, an optimized mycobacterial NDH-2 inhibitor could prove important for the treatment of several different mycobacterial infections.

## Author contributions

YL purified the protein and prepared membrane vesicles, performed assays, acquired cryo-EM images, and calculated and analyzed structures. SAB prepared the NDH-2–3×FLAG *M. smegmatis* strain. GMC provided Compounds 13A and 13C and advised on their use. JLR conceived and supervised the study. YL and JLR wrote the manuscript and prepared the figures with input from the other authors.

## Acknowledgements

We thank Gautier Courbon and Rana Abdelaziz for their assistance in assay design, and Ryan Karimi for assistance with dose-response curve fitting and calculation of IC_50_ values. YL was supported by a Canadian Institutes of Health Research doctoral scholarship and JLR was supported by the Canada Research Chairs program. Cryo-EM data was collected at the Toronto High-Resolution High Throughput cryo-EM facility, supported by the Canada Foundation for Innovation and Ontario Research Fund. Activity assays used infrastructure from the Structural and Biophysical Core Facility at the Hospital for Sick Children. This research was supported by Canadian Institutes of Health Research project grant PJT191893.

## Data availability

Cryo-EM maps are available from the electron microscopy databank with accession codes EMD- 48544, EMD-48546, and atomic models are available from the protein databank with accession codes 9MQY, 9MQZ.

## Materials and Methods

### Generation of an M. smegmatis strain with a C-terminal 3×FLAG tag on NDH-2

*M. smegmatis* strain SABM5, encoding a 3×FLAG sequence 3′ to the gene *MSMEG_3621*, which produces a C-terminal 3×FLAG tag on NDH-2, was generated with the ORBIT method (oligonucleotide-mediated recombineering followed by Bxb1 integrase targeting) (Murphy et al., 2018). Briefly, *M. smegmatis* strain mc^2^155 was transformed with plasmid pKM444 (Addgene #108319). The pKM444-transformed *M. smegmatis* was then transformed with pSAB41 (Guo et al., 2021) and an oligonucleotide (sequence 5′– TGCGCGAACCCGCATCGAGGAGCTCGAGGAGATCGCGGCGGCGGTGCAGGACACCGAGAAAGCCGCGTCCGGTTTGTCTGGTCAACCACCGCGGTCTCAGTGGTGTACGGTACAAACCTAGCCGGTTCGGTCGAGCCGGTTCAGTCGGCAGGCAGCCGGTCACCCTGCCAGGTGGCCAGGCTGCCGCCGCG–3′) that guides integration of pSAB41 into the *M. smegmatis* genome. Hygromycin B (50 μg/mL) was used to select transformants, and correct integration of pSAB41 was confirmed by colony polymerase chain reaction.

### Bacterial culture and isolation of membranes

Cultures of *M. smegmatis* were grown for ∼72 hours at 30 °C with 180 rpm shaking in 2.8 L Fernbach flasks containing 1 L of Middlebrook 7H9 broth supplemented with tryptone (10 g/L), D-glucose (2 g/L), NaCl (0.8 g/L), and 0.05 % (v/v) tween-80. This medium was selected to maximize the mass of cells that could be obtained (Yanofsky et al., 2021). Cells were collected by centrifugation at 6,500 *g* for 20 min and resuspended in 25 mL/L lysis buffer (50 mM HEPES/KOH pH 6.8, 100 mM KH_2_PO_4_, 100 mM NaCl, 5 mM MgCl_2_, 5 mM benzamidine-HCl, 5 mM 6-aminocaproic acid, and 1 mM phenylmethylsulfonyl fluoride) before lysis at > 20 kPa with an Emulsiflex C3 homogenizer (Avestin). Cellular debris was removed by centrifugation at 39,000 *g* for 30 min. Membranes were collected by ultracentrifugation at 200,000 *g* for 60 min before being resuspended in 45 mL of buffer (20 mM HEPES/KOH pH 6.8, 100 mM KH_2_PO_4_, 100 mM NaCl, 5 mM MgCl_2_, 20 % [v/v] glycerol) and frozen at –80 °C.

### Purification of NDH-2

All purification steps were conducted at 4 °C. Frozen membranes were thawed and solubilized with 1 % (w/v) DDM for 60 min. Insoluble material was removed by ultracentrifugation at 200,000 *g* for 50 min. The supernatant was filtered with a 0.45 μm syringe filter (Millipore) and loaded onto a plastic column containing 0.5 mL of M2 anti-FLAG affinity matrix (Sigma) equilibrated with DDM wash buffer (20 mM HEPES/KOH pH 6.8, 100 mM KH_2_PO_4_, 200 mM NaCl, 5 mM MgCl_2_, 15 % [v/v] glycerol, 0.03 % [w/v] DDM). The detergent was exchanged from DDM to LMNG by washing the matrix with 10 column volumes of high LMNG buffer (20 mM HEPES/KOH pH 6.8, 100 mM KH_2_PO_4_, 200 mM NaCl, 5 mM MgCl_2_, 15 % [v/v] glycerol, 0.01 % [w/v] LMNG) followed by two column volumes of low LMNG buffer (20 mM HEPES/KOH pH 6.8, 100 mM KH_2_PO_4_, 200 mM NaCl, 5 mM MgCl_2_, 15 % [v/v] glycerol, 0.001 % [w/v] LMNG). Bound protein was eluted with three column volumes of low LMNG buffer containing 150 μg/mL 3×FLAG peptide. Eluted protein was concentrated with a 30 kDa cut-off concentrator (Millipore) and loaded onto a Superdex 200 increase 10/300 GL equilibrated with LMNG size exclusion buffer (20 mM HEPES/KOH pH 6.8, 100 mM KH_2_PO_4_, 200 mM NaCl, 5 mM MgCl_2_, 0.001 % [w/v] LMNG). Fractions from the Superdex 200 increase column that contained NDH-2 were pooled and concentrated with a 100 kDa cut-off concentrator. Purified NDH-2 was either frozen at –80 °C for enzyme assays or used immediately to prepare cryo-EM grids.

### Comparison of NDH-2 sequence conservation

For comparison of NDH-2 from mycobacteria, amino acid sequences of *M. tuberculosis ndh* (*Rv1854c*) and *ndhA* (*Rv0392c*), *M. leprae* ML2061, *M. avium* MAV_2867 and MAV_4772, *M. abscessus* MAB_2429, and *M. smegmatis* NDH-2 (*Msmeg_3621*) from UniProt (The UniProt Consortium, 2024) were aligned by Multiple Alignment using Fast Fourier Transform (Katoh et al., 2002) (**Supporting Information 1**). For comparison of NDH-2 with known protein structures, amino acid sequences were obtained from UniProt based on the entry used for the structure in the Protein Data Bank: *S. cerevisiae* Ndi1 (PDB: 4G74, UniProt: P32340), *C. thermarum* NDH-2 (PDB: 6BDO, UniProt: F5L3B8), *P. falciparum* NDH-2 (PDB: 5JWC, UniProt: Q8I302), and *S. aureus* NDH-2 (PDB: 5NA1, UniProt: Q2FZV7) (**Supporting Information 2**). The multiple sequence alignments were visualized for publication with ESPript (Robert and Gouet, 2014).

### Preparation of IMVs

IMVs were prepared as described previously (Harden et al., 2024). IMVs for NADH- and succinate-driven acidification assays were from a strain of *M. smegmatis* with a 3×FLAG tag at the C terminus of the QcrB subunit of the Complex III_2_IV_2_ supercomplex (Yanofsky et al., 2021). IMVs for ATP-driven acidification assays were from *M. smegmatis* strain GMC_MSM2 (Guo et al., 2021), where the C-terminal extension of the ATP synthase α subunit is truncated with a 3×FLAG tag, enabling ATP hydrolysis and proton pumping. For the QcrB–3×FLAG strain, cells were collected by centrifugation and resuspended in 50 mM Tris–HCl pH 7.5 (at room temperature), 150 mM NaCl, 5 mM MgSO_4_, 5 mM benzamidine HCl, and 5 mM 6-aminocaproic acid. Cells were lysed with four passes through an Emulsiflex C3 homogenizer and cellular debris removed through centrifugation at 39,000 *g* for 30 min. Membranes were collected by ultracentrifugation at 200,000 *g* for 60 min and resuspended at 2.5 mL/L culture in IMV resuspension buffer (50 mM Tris–HCl pH 7.5 at room temperature, 150 mM NaCl, 5 mM MgSO_4_, 5 mM benzamidine HCl, 5 mM 6-aminocaproic acid, and 20 % [v/v] glycerol). For preparation of IMVs from *M. smegmatis* strain GMC_MSM2, membranes were collected as with QcrB-3×FLAG but were resuspended in IMV resuspension buffer at 2.7 mL/L culture. IMVs were frozen and stored at –80 °C.

### Activity assays

To measure NADH:quinone oxidoreductase activity, purified NDH-2 was incubated with 1 mM decylubiquinone in assay buffer (10 mM MES pH 6, 25 mM NaCl, 5 mM CaCl_2_, 0.05% [w/v] CHAPS, 0.05% [w/v] azolectin) in a 96 well clear flat bottom micro test plate (Sarstedt). The reaction was started by addition of 200 μM NADH and absorbance at 340 nm, which is proportional to the concentration of NADH, was measured with a SpectraMax i3x Multi-Mode Microplate Reader (Molecular Devices) at 37 °C.

To measure the effects of inhibitors on IMV acidification, fluorescence from IMV solutions with ACMA in a BRANDplates black U-bottom 96 well microtitration plate (BRANDTECH Scientific) was measured with a BioTek Synergy Neo2 plate reader (Agilent) at 25 °C. The excitation wavelength was set to 410 nm, the emission wavelength to 480 nm, and the fluorescence gain was set to 90. The necessary dilution of IMV solutions to maximize signal was determined for each batch of IMVs before the experiment. Assays were initiated by adding NADH, succinate, or ATP to 10 µL of diluted IMV solution in a total reaction volume of 160 µL, so that each well contained 10 mM HEPES–KOH pH 7.5, 100 mM KCl, 5 mM MgCl_2_, 2 % (v/v) DMSO and 3.125 μM ACMA, as well as either 1 mM NADH, 5 mM sodium succinate, or 8 mM ATP buffered with 16 mM Tris (pH unadjusted). When used, Compounds **13A** and **13C** (Shirude et al., 2012) were added from 1 mM stocks in DMSO without changing the final concentration of DMSO in the well, while Compound **1** (Molport) was added from 10 mM stocks in DMSO without changing the final concentration of DMSO in the well. IMV acidification was quantified by measuring the fluorescence recovery following addition of 3.2 μL of 100 μM nigericin in 1% ethanol.

### Cryo-EM specimen preparation and data collection

Holey gold foils with 2 μm holes in a square lattice were nanofabricated on 400 mesh copper– rhodium mesh grids as described previously (Marr et al., 2014). Purified NDH-2 was concentrated to ∼2 mg/mL as measured by bicinchoninic acid assay (Pierce). For the inhibitor-free sample, grids were glow-discharged for 15 s in air and 1.5 μL of sample was applied in the environmentally controlled chamber of an EM GP2 Plunge Freezer (Leica Microsystems), held at 4 °C and with 90–95 % nominal relative humidity. Grids were blotted for 0.5 seconds and plunge frozen in liquid ethane. For the inhibitor-bound sample, 0.2 μL of 1.1 mM Compound **1** was added to 8.6 μL of concentrated NDH-2 to reach a final inhibitor concentration of 25 μM. The sample was incubated at room temperature for 5 min before removing insoluble material by centrifugation at 2000 *g* for 10 s, and then grids were frozen as described above.

Both specimen screening and high-resolution data collection were automated with the EPU software (ThermoFisher Scientific). All Samples were screened with a Glacios 2 electron microscope (ThermoFisher Scientific) operating at 200 kV and equipped with a Falcon 4i camera. Images were collected at 92,000× nominal magnification in counting mode, corresponding to a calibrated pixel size of 1.5 Å, with a total exposure of ∼40 e^−^/Å^2^. For the high-resolution inhibitor-free sample, 14,783 movies were collected in Electron Event Representation (EER) format (Guo et al., 2020) with a Titan Krios G3 electron microscope (ThermoFisher Scientific) operating at 300 kV and equipped with a Falcon 4i direct electron detector. The nominal magnification was 120,000×, corresponding to a calibrated pixel size of 0.64 Å. A total exposure of ∼70 e^−^/Å^2^ was used. For the inhibitor-bound sample, 9,155 movies in EER format were collected with a 300 kV Titan Krios G3 electron microscope equipped with a Selectris X energy filter (ThemoFisher Scientific) with a slit width of 10 eV and a Falcon 4i direct electron detector. The nominal magnification was 165,000×, corresponding to a calibrated pixel size of 0.73 Å, and a total exposure of ∼70 e^−^/Å^2^ was used.

### Cryo-EM image analysis

Data quality during automated collection was assessed with cryoSPARC Live. All image analysis was done with cryoSPARC v4.6.2 (Punjani et al., 2017). Exposure fractions were aligned (Rubinstein and Brubaker, 2015), and contrast transfer function (CTF) parameters estimated in patches. All map refinement was done without symmetry until the final refinement, where C2 symmetry was enforced. For the inhibitor-free sample, templates for particle selection were projections of a low-resolution 3D map calculated from a dataset obtained during specimen screening. Selected particle images were extracted from a subset of 4000 micrographs with a box size of 360×360 pixels, Fourier cropped to 180×180 pixels (1.28 Å/pixel), and subjected to 2D classification. Particle images from classes where the class average showed clear secondary structure features were used to generate two separate Topaz picking models (Bepler et al., 2019), one for the “top” view and one for the “side” views of NDH-2. Topaz models were refined with iterative rounds of particle extraction, 2D classification, and retraining. Following the final round of Topaz picking, particle images selected with either model were combined, duplicates removed with a minimum separation distance of 70 Å, and 3,118,052 particle images were extracted with a box size of 384×384 pixels and Fourier cropped to 96×96 pixels (2.56 Å/pixel). The dataset of particle images was cleaned with one round of 2D classification, and initial 3D maps were calculated by *ab initio* reconstruction from particle images from selected 2D class averages.

Micrographs where no particle images were extracted were removed and 1,093,606 particle images were extracted from 12,245 micrographs with a box size of 384×384 (0.64 Å/pixel). This dataset was cleaned with multiple rounds of heterogeneous refinement before a high-resolution map was calculated by non-uniform refinement (Punjani et al., 2020). Per-particle CTF parameters were measured, and the particles were subjected to a final round of 3D classification without alignment to remove particle images that contributed noise to the map. The final 3D map from 59,561 particle images was calculated using local refinement with a mask that excluded the detergent micelle.

For the Compound **1**-bound dataset, initial particle selection was done on a subset of 800 micrographs using the final Topaz model from the inhibitor-free dataset. Extracted particle images were cleaned with 2D classification and particle images from selected 2D classes were used to train separate Topaz models for “top” and “side” views of NDH-2. Topaz was then used to select particle images from the full dataset. Following removal of duplicate particles, a total of 1,066,096 particle images were extracted with a box size of 384×384 pixels and Fourier cropped to 96×96 pixels (2.56 Å/pixel). These particle images were used in heterogeneous refinement with four reference maps: a lowpass filtered map of inhibitor-free NDH-2, and three low-quality maps calculated *ab initio* from the particle images. Only particle images that were assigned to the lowpass filtered map of inhibitor-free NDH-2 in heterogenous refinement were selected for *ab initio* reconstruction followed by heterogeneous refinement. 181,889 particle images corresponding to the selected class following heterogeneous refinement were re-extracted with a box size of 384×384 pixels (0.73 Å/pixel) and cleaned further with multiple rounds of *ab initio* reconstruction and heterogeneous refinement. A final round of cleaning was done by 3D classification without alignment to remove particle images that contributed noise to the map. The final 3D map from particle images was calculated using local refinement with a mask that excluded the detergent micelle.

### Model building and refinement

For inhibitor-free NDH-2, the starting model of the monomer was obtained from the AlphaFoldDB protein structure database (Jumper et al., 2021; Varadi et al., 2022). Two copies of the NDH-2 monomer were fit as rigid bodies into the cryo-EM map with UCSF Chimera (Pettersen et al., 2004). Optimization of model-to-map fit and molecular dihedral angles, as well as incorporation and fitting of the model of the FAD cofactor was done with Coot v0.9.6 (Emsley and Cowtan, 2004) and ISOLDE v1.8 (Croll, 2018). Model parameters were refined with PHENIX v1.19.2 (Liebschner et al., 2019). For Compound **1**-bound NDH-2, the refined inhibitor-free NDH-2 atomic model was used as a starting model. A 3D model of Compound **1** was generated with *phenix.elbow* from the SMILES (Simplified Molecular Input Line Entry System) string. Fitting of the Compound **1** model into the cryo-EM density was done with Coot and the model was optimized and refined as done for the inhibitor-free NDH-2 model. Figures were rendered with UCSF Chimera and UCSF ChimeraX (Goddard et al., 2018).

## Notes

### Competing Interest Statement

The authors have declared no competing interest.

