## Supporting Information 1 for "Structure of mycobacterial NDH-2 bound to a 2-mercapto-quinazolinone inhibitor"

Supporting information 1: Alignment of NDH-2 amino acid sequences from mycobacterial species.

```
                   1                                   10        20        30
smegmatis          MS.HPGATAS.................DRHKVVIIGSGFGGLTAAKTLKR....ADVDVK
leprae             MNAQPAATAAQD...............RRHQVVIIGSGFGGLNAAKKLKR....ANVDIK
tuberculosis_ndh   MSPQQEPTAQPP...............RRHRVVIIGSGFGGLNAAKKLKR....ADVDIK
avium_2867         MSPHSGSTAGPE...............RRHQVVIIGSGFGGLNAAKKLKH....ANVDIK
abcessus_2429      MSTTPDAAGAAAPVAANPQPKIQDALTPKHRVVIIGSGFGGLTAAKTLKR....ANADVT
tuberculosis_ndhA  MTLSSGEPSAVG...............GRHRVVIIGSGFGGLNAAKALKR....ADVDIT
avium_4772         MS..........................RHRVVIIGSGFGGLTAAKALKRVPKGTQVDIT

                     40        50        60        70        80        90
smegmatis          LIARTTHHLFQPLLYQVATGIISEGEIAPATRVILRKQKNAQVLLGDVTHIDLENKTVDS
leprae             LIARTTHHLFQPLLYQVATGIISEGEIAPPTRVVLRKQRNIQVLLGNVTHIDLANQCVVS
tuberculosis_ndh   LIARTTHHLFQPLLYQVATGIISEGEIAPPTRVVLRKQRNVQVLLGNVTHIDLAGQCVVS
avium_2867         LIARTTHHLFQPLLYQVATGIVSEGDIAPPTRVVLRRQRNVQVLLGDVTHIDLAGKFVVS
abcessus_2429      LIARTTHHLFQPLLYQVATGIISEGEIAPPTRQILKDQDNAQVVLGDVTSIDLEKQVVHS
tuberculosis_ndhA  LISKTTTHLFQPLLYQVATGILSEGDIAPTTRLILRRQKNVRVLLGEVNAIDLKAQTVTS
avium_4772         LISKTTTHLFQPLLYQVATGILSEGDIAPTTRLILRKQKNVRVLWGDVSAIDLTARTVTS

                   100       110       120       130       140       150
smegmatis          VLLGHTYSTPYDSLIIAAGAGQSYFGNDHFAEFAPGMKSIDDALELRGRILGAFEQAERS
leprae             ELLGHTYETLYDSLIVAAGAGQSYFGNDQFAEFAPGMKSIDDALELRGRILSAFEQAERS
tuberculosis_ndh   ELLGHTYQTPYDSLIVAAGAGQSYFGNDHFAEFAPGMKSIDDALELRGRILSAFEQAERS
avium_2867         DLLGHTYETPYDSLIVAAGAGQSYFGNDHFAEFAPGMKSIDDALEVRGRILSAFEQAERS
abcessus_2429      SLLGHDYSTPYDSLIVAAGAGQSYFGNDHFAEWAPGMKSIDDALELRGRILGAFEQAERS
tuberculosis_ndhA  KLMDMTTVTPYDSLIVAAGAQQSYFGNDEFATFAPGMKTIDDALELRGRILGAFEAAEVS
avium_4772         HLMGMDTVTPYDSLIVAAGAQQSYFGHDEYAAFAPGMKSVDDALELRGRILGAFEAAEVA

                   160       170       180       190       200       210
smegmatis          SDPVRRAKLLTFTVVGAGPTGVEMAGQIAELADQTLRGSFRHIDPTEARVILLDAAPAVL
leprae             NDPERREKLLTFTVVGAGPTGVEMAGQIAELADHTLKGAFRRIDSTKARVVLLDAAPAVL
tuberculosis_ndh   SDPERRAKLLTFTVVGAGPTGVEMAGQIAELAEHTLKGAFRHIDSTKARVILLDAAPAVL
avium_2867         RDPERRAKLLTFTVIGAGPTGVEMAGQIAELATYTLKGSFRHIDPTKARVILLDAAPAVL
abcessus_2429      SDPVRRRKLMTFTVVGAGPTGVEMAGQIAELANETLKGTFRHIDPTDARVILLDAAPAVL
tuberculosis_ndhA  TDHAERERRLTFVVVGAGPTGVEVAGQIVELAERTLAGAFRTITPSECRVILLDAAPAVL
avium_4772         TDPAERRRRLTFVVVGAGPTGVELAGEIVQLAERTLAGAFRTITPSECRVILLDAAPAVL

                   220       230       240       250       260       270
smegmatis          PPMGEKLGKKARARLEKMGVEVQLGAMVTDVDRNGITVKDSDGTIRRIESACKVWSAGVS
leprae             PPMGEKLGQRAAARLQKMGVEIQLSAMVTDVDRNGITVQDSDGTVRRIESACKVWSAGVS
tuberculosis_ndh   PPMGAKLGQRAAARLQKLGVEIQLGAMVTDVDRNGITVKDSDGTVRRIESACKVWSAGVS
avium_2867         PPFGDKLGKRAADRLEKMGVEIQLGAMVTDVDRNGITVKDSDGTVRRIESACKVWSAGVS
abcessus_2429      PPFGPKLGDKARKRLEKLGVEIQLSAMVTDVDRNGLTVKHADGSIERIESWCKVWSAGVS
tuberculosis_ndhA  PPMGPKLGLKAQRRLEKMDVEVQLNAMVTAVDYKGITIKEKDGGERRIECACKVWAAGVA
avium_4772         PPMGPKLGLKAQRRLQKMDVEVQLNAMVTAVDYMGITVKEKDGAERRIECACKVWAAGVQ

                   280       290       300       310       320       330
smegmatis          ASPLGKDLAEQS.GVELDRAGRVKVQPDLTLPGHPNVFVVGDMAAVEGVPGVAQGAIQGG
leprae             ASRLGRDLAEQS.LVELDWAGRVKVLPDLSIPGYRNVFVVGDMAAIDGVPGVAQGAIQGA
tuberculosis_ndh   ASRLGRDLAEQS.RVELDRAGRVQVLPDLSIPGYPNVFVVGDMAAVEGVPGVAQGAIQGA
avium_2867         ASPLGRDLAEQS.TVELDRAGRVKVLPDLSIPGHPNVFVIGDLAAVEGVPGVAQGAIQGA
abcessus_2429      ASPLGKNLAEQS.GVELDRAGRVKVGPDLSIPGHPNVFVVGDMMAVDGVPGVAQGAIQGG
tuberculosis_ndhA  ASPLGKMIAEGSDGTEIDRAGRVIVEPDLTVKGHPNVFVVGDLMFVPGVPGVAQGAIQGA
avium_4772         ASALGAMIAEQSDGTETDRAGRVIVEPDLTVKGHPYVFVVGDLMSVPGVPGMAQGAIQGA

                   340       350       360       370       380       390
smegmatis          RYAAKIIKREVSGT.SPKIRTPFEYFDKGSMATVSRFSAVAKVGPVEFAGFFAWLCWLVL
leprae             KYVANNIKAELGEA.NPAEREPFQYFDKGSMATVSRFSAVAKIGPLEFSGFIAWLMWLVL
tuberculosis_ndh   KYVASTIKAELAGA.NPAEREPFQYFDKGSMATVSRFSAVAKIGPVEFSGFIAWLIWLVL
avium_2867         KYVANTIKAELGGA.DPAEREPFQYFDKGSMATVSRFSAVAKIGPLEFSGLFAWFAWLVL
abcessus_2429      RYAAKQIKAELKGR.SPAERAPFKFFDKGSMATVSRYSAVVKIGKIEFSGFIAWLSWLAL
tuberculosis_ndhA  RYATTVIKHMVKGNDDPANRKPFHYFNKGSMATISRHSAVAQVGKLEFAGYFAWLAWLVL
avium_4772         KYAAGIIKQAVKGQDNPAERKPFRYVNKGSMALISRYSAVAQVGRVEFAGFIAWLAWLLL

                     400       410       420       430       440       450
smegmatis          HLVYLVGFKTKIVTLLSWGVTFLSTKRGQLTITEQQAYAR...TRIEELEE...IAAAVQ
leprae             HLMYLIGFKTKITTLLSWTVTFLSTRRGQLTITDQQAFAR...TRLEQLAE...LTAESQ
tuberculosis_ndh   HLAYLIGFKTKITTLLSWTVTFLSTRRGQLTITDQQAFAR...TRLEQLAE...LAAEAQ
avium_2867         HLVYLVGFKTKVSTLLSWTVTFLSTRRGQLTITEQQAFAR...TRLEQLAV...LAAETK
abcessus_2429      HLVYLVGFKNRLVTLILWSIAFVTRSRGQMAITEQQAWAR...TRLDQLEE...LVAEKN
tuberculosis_ndhA  HLVYLVGYRNRIAALFAWGISFMGRARGQMAITSQMIYARLVMTLMEQQAQ.GALAAAEQ
avium_4772         HLYYLIGHRNRLAAMFAWGISFLGRTRGQMAITSQMIYARPVVEWLERRAQDGTLAAAEH

smegmatis          DTEKAAS.........
leprae             STAAANRATMR...AS
tuberculosis_ndh   GSAASAKV......AS
avium_2867         RPAARR........AS
abcessus_2429      GDEEPTTQPDKEKAAS
tuberculosis_ndhA  AEHAEQEA......AG
avium_4772         VDSAEKQA......G.
```
