## Supporting Information 2 for "Structure of mycobacterial NDH-2 bound to a 2-mercapto-quinazolinone inhibitor"

Supporting information 2: Alignment of NDH-2 amino acid sequences from species where experimental structures have been determined.

```
               1                                                10
M_smegmatis    MS...............................HPGATA...........SDRHKVVIIG
C_thermarum    MS..................................................KPSIVILG
S_aureus       MA................................................QDRKKVLVLG
S_cerevisiae   MLSKNLYSNKRLLTSTNTLVRFASTRSTGVENSGAGPTSFKTMKVIDPQHSDKPNVLILG
P_falciparum   MLVKF.....RKCGQANIFRSISNVRKI....YNVAKNNLKNNKDI....ERKEKIIILG

               20        30        40        50        60        70
M_smegmatis    SGFGGLTAA....KTLKRADVDVKLIARTTHHLFQPLLYQVATGIISEGEIAPATRVILR
C_thermarum    AGYGGIVAALGLQKRLNYNEADITLVNKNDYHYITTELHQPAAGTMHHDQARVGIKELID
S_aureus       AGYAGLQTVTKLQKAISTEEAEITLINKNEYHYEATWLHEASAGTLNYEDVLYPVESVLK
S_cerevisiae   SGWGAISFL....KHIDTKKYNVSIISPRSYFLFTPLLPSAPVGTVDEKSIIEPIVNFAL
P_falciparum   SGWGGFNFL....LNIDFKKYDVTLISPRNYFTFTPLLPCLCSGTLSVNVCTESIRNFLR

                      80        90        100                       110
M_smegmatis    KQK..NAQVLLGDVTHIDLENKTVDSVLL..........GHT.......YSTPYDSLIIA
C_thermarum    EKK...IKFVKDTVVAIDREQQKVTLQN.....................GELHYDYLVVG
S_aureus       KDK...VNFVQAEVTKIDRDAKKVETNQ.....................GIYDFDILVVA
S_cerevisiae   KKKG.NVTYYEAEATSINPDRNTVTIKSLSAVSQLYQPENHLGLHQAEPAEIKYDYLISA
P_falciparum   KKNGYCGNYLQLECTDVFYEDKYINCIDI.........ENNK.......VKLFYDYLIIA

                  120       130       140       150       160       170
M_smegmatis    AGAGQSYFGNDHFAEFAPGMKSIDDALELRGRILGAFEQAE..RSSDPVRRAKLLTFTVV
C_thermarum    LGSEPETFGIEGLREHAFSINSINSVRIIRQHIEYQFAKF....AAEPERTDY.LTIVVG
S_aureus       LGFVSETFGIEGMKDHAFQIENVITARELSRHIEDKFANY....AASKEKDDNDISILVG
S_cerevisiae   VGAEPNTFGIPGVTDYGHFLKEIPNSLEIRRTFAANLEKANLLPKGDPERRRL.LSIVVV
P_falciparum   VGAKTNTFNINGVDKYAYFVKDIDDALKIRKKFLDILEKCTLPNISNEEKKKM.LHVAVV

                    180       190       200       210       220       230
M_smegmatis    GAGPTGVEMAGQIAELADQTLRGSFRHIDPTEARVILLDAAPAVLPPMGEKLGKKARARL
C_thermarum    GAGFTGIEFVGELADRMPELC.AEYD.VDPKLVRIINVEAAPTVLPGFDPALVNYAMDVL
S_aureus       GAGFTGVEFLGELTDRIPELC.SKYG.VDQNKVKITCVEAAPKMLPMFSEELVNHAVSYL
S_cerevisiae   GGGPTGVEAAGELQDYVHQDLRKFLP.ALAEEVQIHLVEALPIVLNMFEKKLSSYAQSHL
P_falciparum   GGGPTGVEVTAEFADFINKEVKINYK.DIFNFISISIIEGGNNLLPTFTQNISDFTKENF

                    240       250       260       270       280
M_smegmatis    EKMGVEVQLGAMVTDVDRNGITVKDS..DG..TIRRIESACKVWSAGVSASPLGKDLAEQ
C_thermarum    GGKGVEFKIGTPIKRCTPEGVVIEV...DG..EEEEIKAATVVWTGGVRGNSIVEKSGFE
S_aureus       EDRGVEFKIATPIVACNEKGFVVEV...DG..EKQQLNAGTSVWAAGVRGSKLMEESFEG
S_cerevisiae   ENTSIKVHLRTAVAKVEEKQLLAKTKHEDGKITEETIPYGTLIWATGNKARPVITDLFKK
P_falciparum   HNLNINVLTNYYVIDVDKHSFHIQSSLNKN..EKKKLSYGLLIWASGLAQTTLIQKFLKT

             290       300       310       320
M_smegmatis    SGVELDRAGRVKVQPDLTLPG..HPNVFVVGDMAAV........................
C_thermarum    T.....MRGRIKVDPYLRAPG..HENIFIVGDCALIIN......................
S_aureus       V.....KRGRIVTKQDLTING..YDNIFVIGDCSAFIP......................
S_cerevisiae   IPEQNSSKRGLAVNDFLQVKG..SNNIFAIGDNAF.........................
P_falciparum   IPVQA.NNAILKVDEKLRVIGIPSNNIYAIGDCKKIQPKLLHEHTNEIIKILTGNKLTSE

                                                                 330
M_smegmatis    ..........EGVP.....................................GVAQGAIQG
C_thermarum    ......EENNRPYP.....................................PTAQIAIQH
S_aureus       ......AGEERPLP.....................................TTAQIAMQQ
S_cerevisiae   ..........AGLP.....................................PTAQVAHQE
P_falciparum   ALKLKQSELTKTFPQLSISKWDYEKNKKGEMTPQQFHDYLFEIDKNYKSPTPTAQNAKQE

                 340       350                 360       370
M_smegmatis    GRYAAK.........IIKREVSGTSPKIRT........PFEYFDKGSMATVSRFSAVAKV
C_thermarum    GENVAA.........NLAALIRGGSMT...........PFKPHIRGTVASLGRNDAIGIV
S_aureus       GESVAK.........NIKRILNGESTE...........EFEYVDRGTVCSLGSHDGVGMV
S_cerevisiae   AEYLAKNFDKMAQIPNFQKNLSSRKDKIDLLFEENNFKPFKYNDLGALAYLGSERAIATI
P_falciparum   AYYLSN.........VFNNFIHTNQKF.......NIPSFIEKWKGSLAYIGNHQVVADL

                380       390       400       410       420       430
M_smegmatis    ..GPVEF...AGFFAWLCWLVLHLVYLVGFKTKIVTLLSWGVTFLSTKRGQLTITEQQAY
C_thermarum    .GGRKVY....GHAASWLKKLIDMRYLYL........IGG..LSLVLKKGRF........
S_aureus       .FGKPIA....GKKAAFMKKVIDTRAVFK........IGG..IGLAFKKGKF........
S_cerevisiae   RSGKRTFYTGGGLMTFYLWRILYLSMILSARSRLKVFFDW..IKLAFFKRDFFKG.....
P_falciparum   ....PYYELKGGRFSSTFWKVVYIQLLLSWKSRFHFFIDF..IKTKWYGRPFIK......

                  440       450
M_smegmatis    ARTRIEELEEIAAAVQDTEKAAS
C_thermarum    ......................
S_aureus       ......................
S_cerevisiae   ......................L
P_falciparum   ......................
```
