## Supplementary material for "Structure of mycobacterial NDH-2 bound to a 2-mercapto-quinazolinone inhibitor": SI_Figures

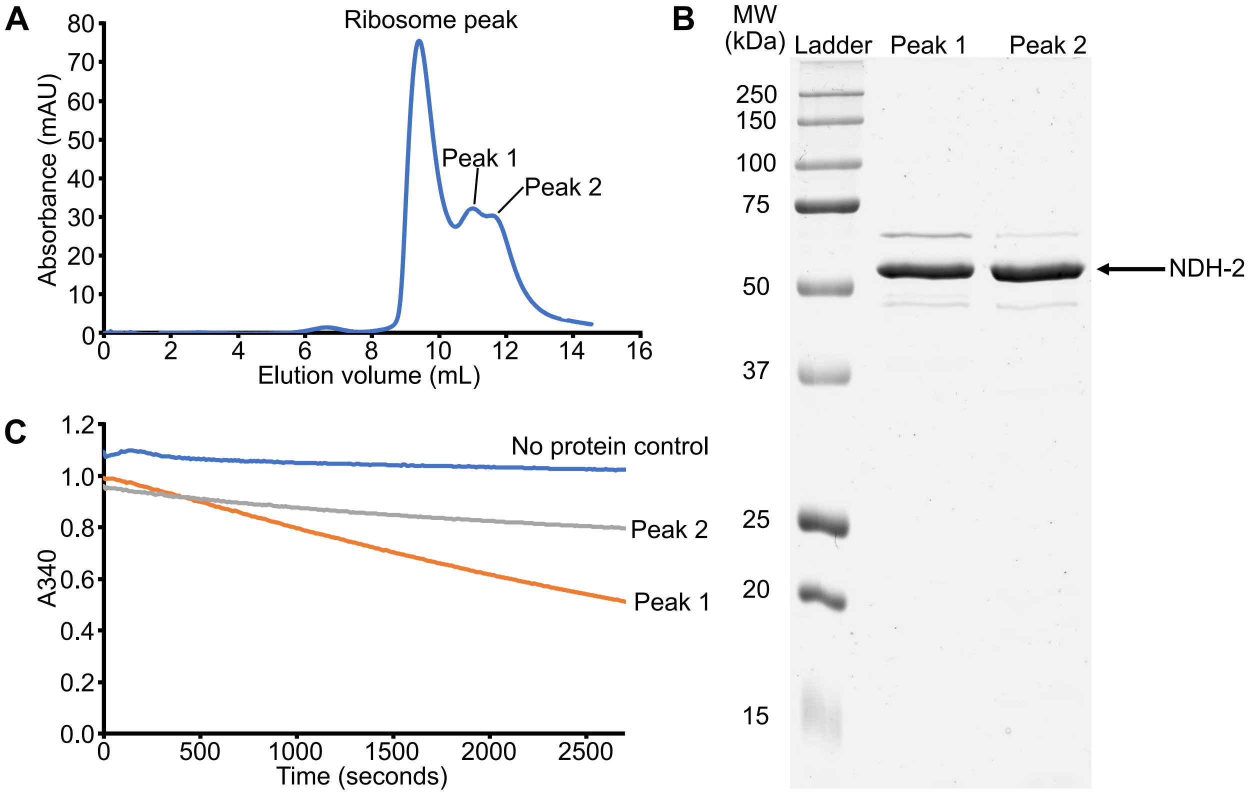


**Supplementary Figure 1: Purification of *M. smegmatis* NDH-2. A)** Size exclusion chromatography (SEC) chromatogram for NDH-2 following anti-FLAG affinity purification. **B)** Corresponding SDS-PAGE gel of SEC-purified NDH-2. **C)** NADH oxidation by purified NDH-2 in the presence of decylubiquinone as an electron acceptor. The concentration of NDH-2 in the assay is not known.

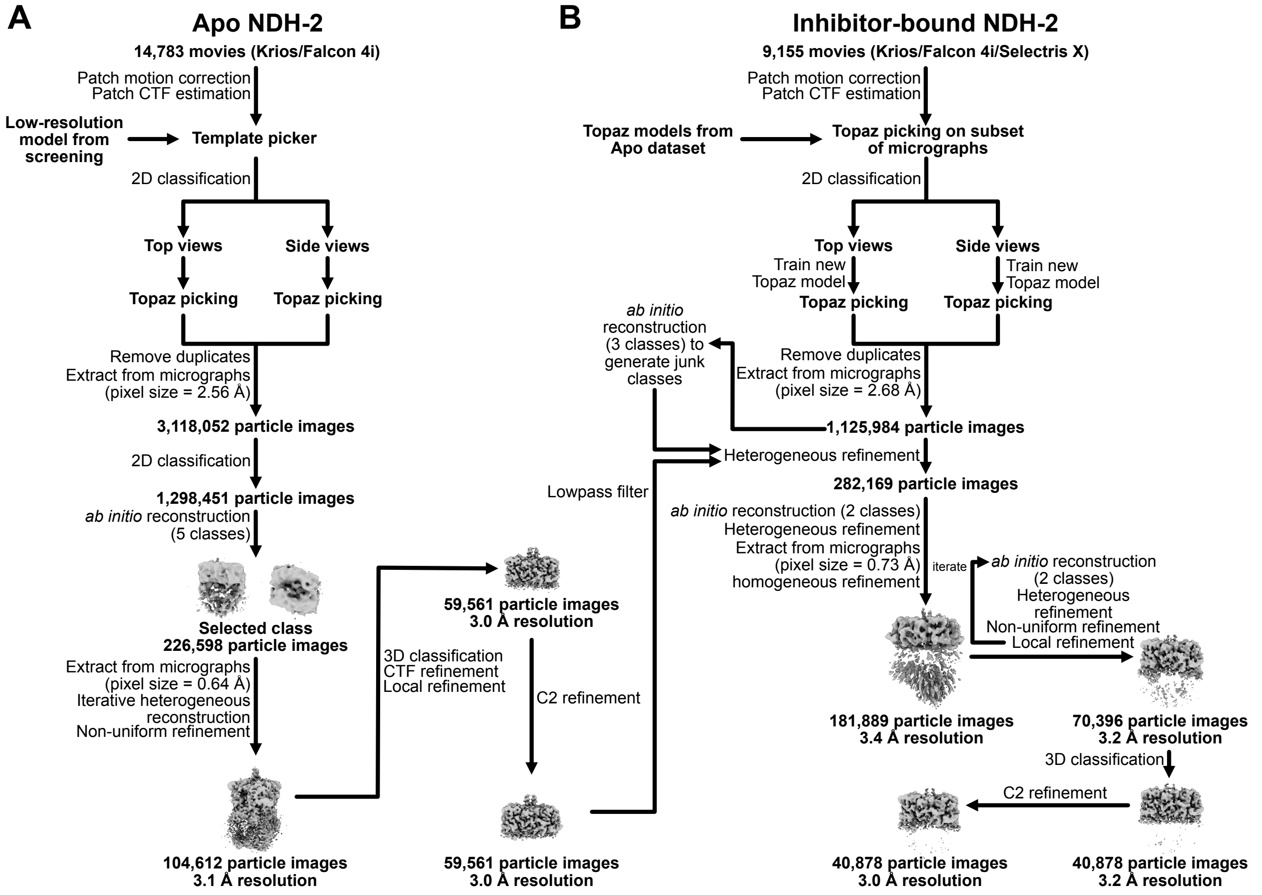


**Supplementary Figure 2: NDH-2 cryo-EM workflow and data processing. A)** Workflow for inhibitor-free NDH-2. **B)** Workflow for Compound **1**-bound NDH-2.

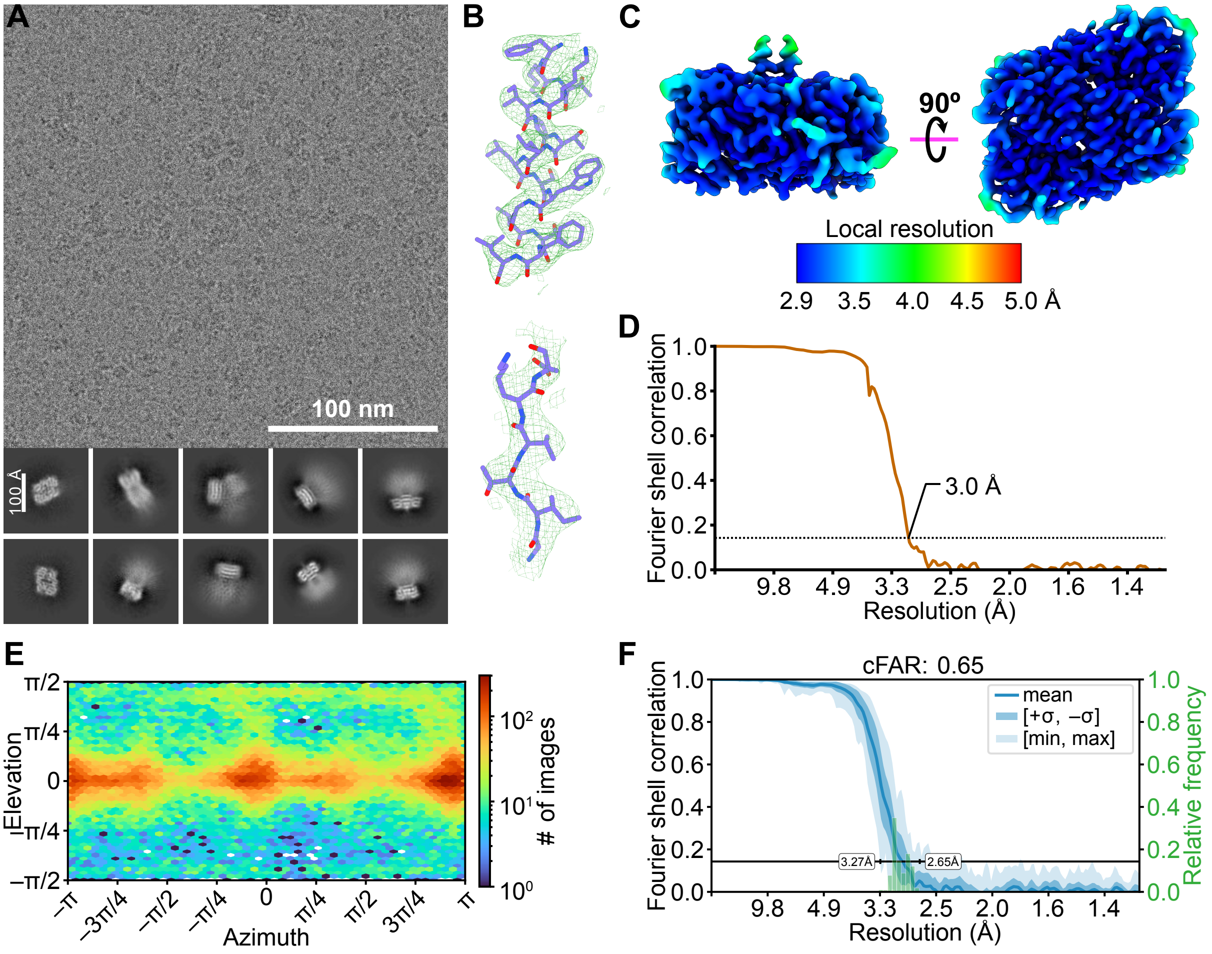


**Supplementary Figure 3: Cryo-EM map validation for inhibitor-free NDH-2. A)** Example cryo-EM micrograph (*top*) and two-dimensional class average images (*bottom*). **B)** *E*xample of model-in-map fit for an α helix (*top*) and a β strand (*bottom*) of the map refined with C2 symmetry. **C)** Local resolution in the cryo-EM map of NDH-2 with C2 symmetry. **D)** Fourier shell correlation (FSC) curve for the cryo-EM map with C2 symmetry following a gold-standard refinement and correction for the effects of masking. **E)** Viewing direction distribution plot of particle images for the map with C2 symmetry. **F)** Conical FSC (cFSC) curve for the cryo-EM map with C2 symmetry.

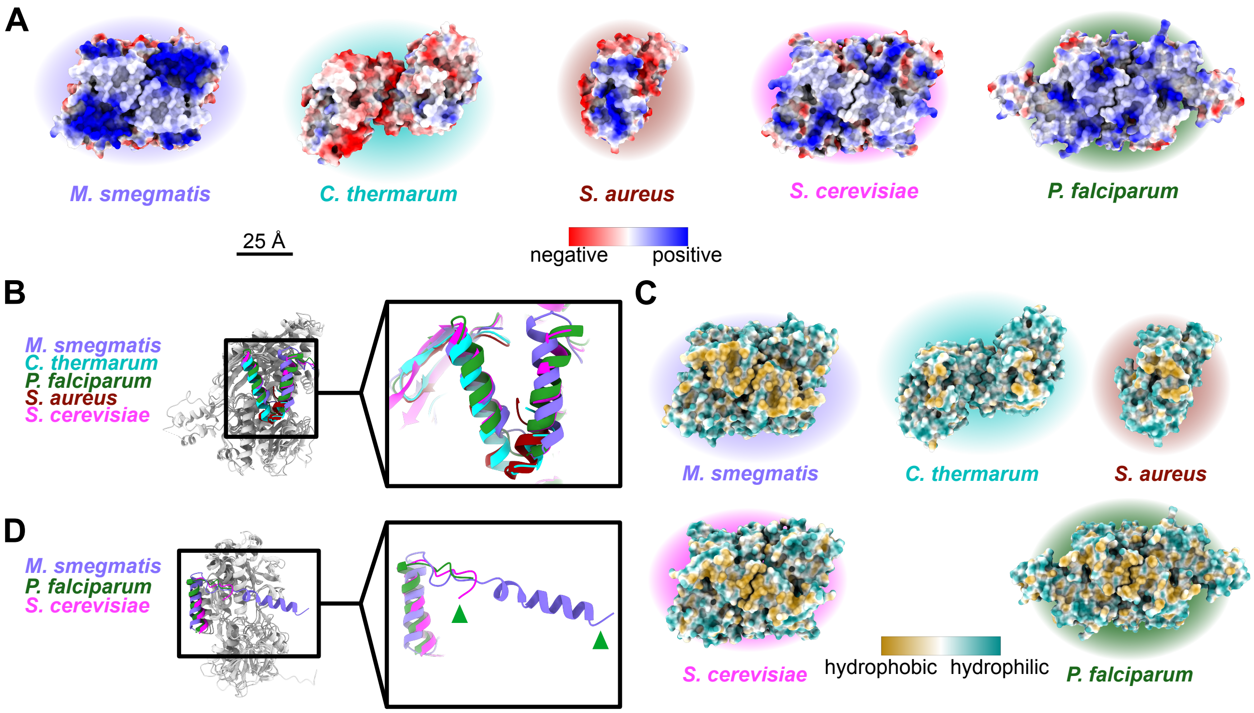


**Supplementary Figure 4: Comparison of NDH-2 structures from different species. A)** Comparison of surface electrostatics. **B)** Comparison of membrane anchoring α helices for experimental structures of NDH-2 from different species. **C)** Comparison of membrane face surface hydrophobicity. **D)** Comparison of the C-terminal α helix from mycobacterial NDH-2 and eukaryotic NDH-2s.

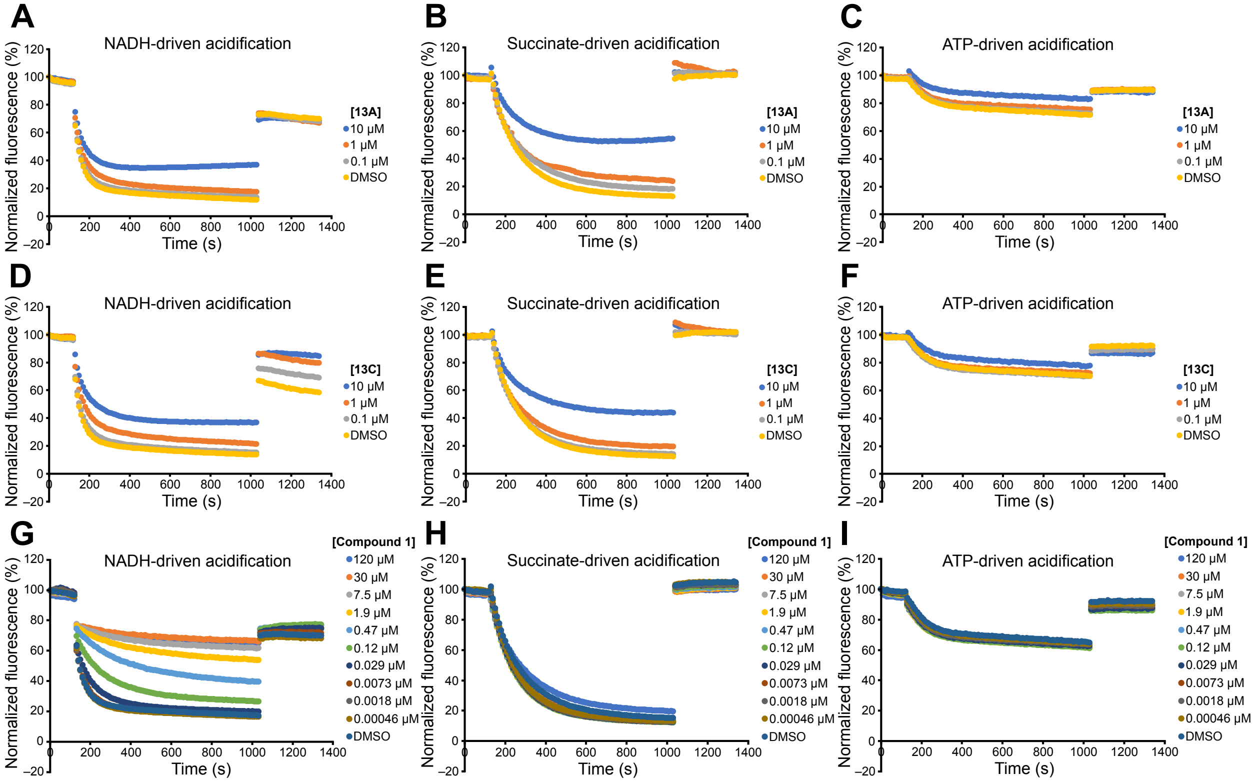


**Supplementary Figure 5: Sample IMV acidification assays measured by ACMA fluorescence quenching. A)** NADH-driven acidification at different concentrations of Compound **13A**. **B)** Succinate-driven acidification at different concentrations of Compound **13A**. **C)** ATP-driven acidification at different concentrations of Compound **13A**. **D)** NADH-driven acidification at different concentrations of Compound **13C**. **E)** Succinate-driven acidification at different concentrations of Compound **13C**. **F)** ATP-driven acidification at different concentrations of Compound **13C**. **G)** NADH-driven acidification at different concentrations of Compound **1**. **H)** Succinate-driven acidification at different concentrations of Compound **1**. **I)** ATP-driven acidification at different concentrations of Compound **1**. Each curve is the average of three experiments.

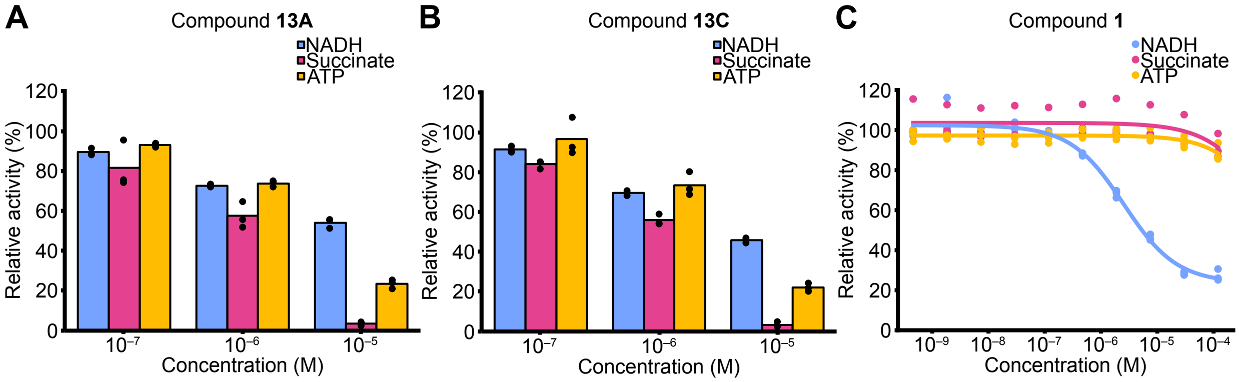


**Supplementary Figure 6: Replicate of assays from independent protein purifications. A)** Replicate of NADH- (*blue*), succinate- (*magenta*), and ATP-driven (*yellow*) IMV acidification at different concentration of quinolinyl pyrimidine Compound **13A**. **B)** Replicate of NADH- (*blue*), succinate- (*magenta*), and ATP-driven (*yellow*) IMV acidification at different concentration of quinolinyl pyrimidine Compound **13C**. **C)** Replicate of dose-response curves for NADH- (*blue*), succinate- (*magenta*), and ATP-driven (*yellow*) IMV acidification at different concentrations of 2-mercapto-quinazolinone Compound **1**.

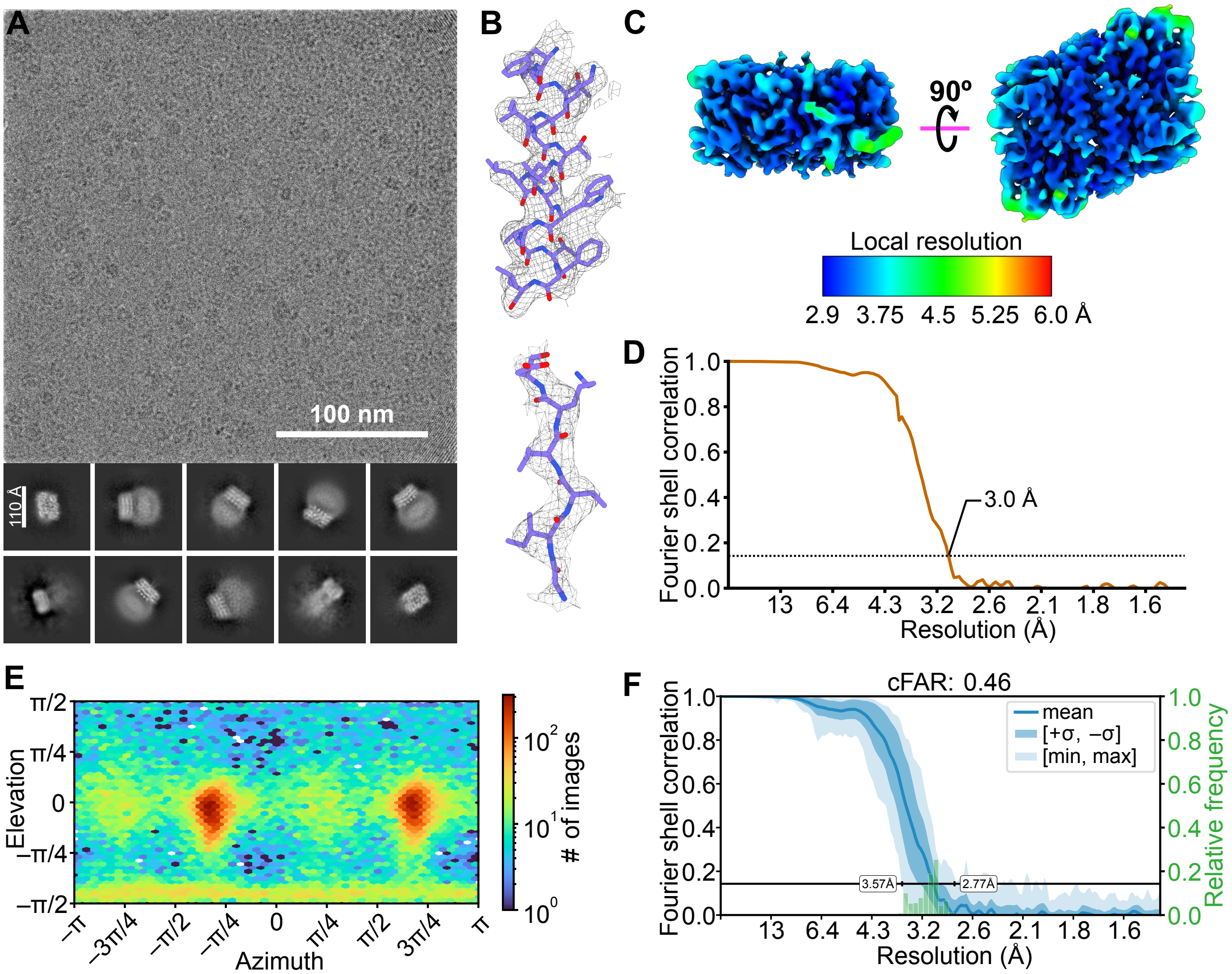


**Supplementary Figure 7: Cryo-EM map validation for NDH-2 bound to Compound 1. A)** Example cryo-EM micrograph (*top*) and two-dimensional class average images (*bottom*). **B)** Example of model-in-map fit for an α helix (*top*) and a β strand (*bottom*) of the map refined with C2 symmetry. **C)** Local resolution in the cryo-EM map of NDH-2 with C2 symmetry. **D)** FSC curve for the cryo-EM map with C2 symmetry following a gold-standard refinement and correction for the effects of masking. **E)** Viewing direction distribution plot of particle images for the map with C2 symmetry. **F)** cFSC curve for the cryo-EM map with C2 symmetry.

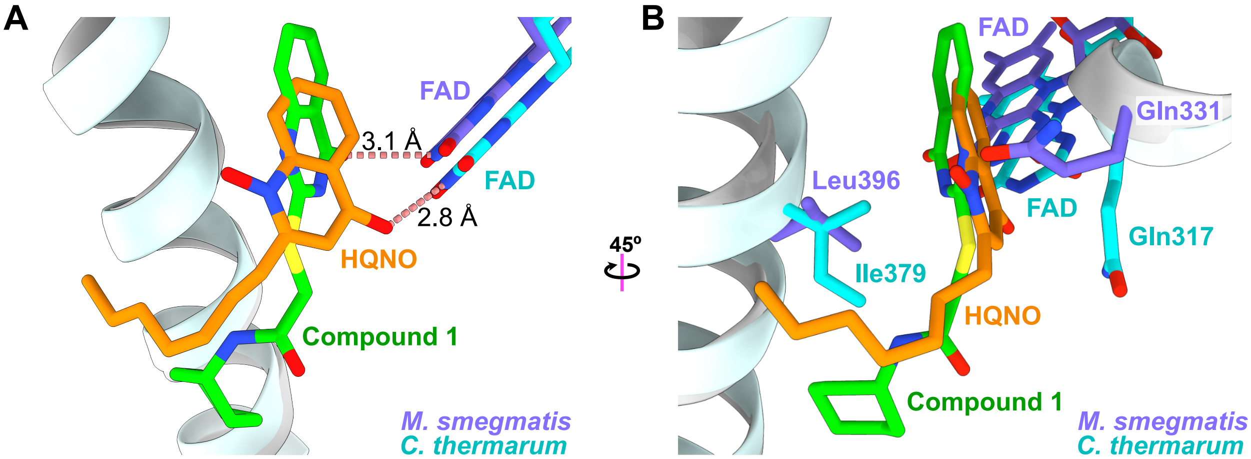


**Supplementary Figure 8: Comparison of Compound 1 and HQNO binding sites. A)** Comparison of position along the quinone-binding site of Compound **1** (*green*) in *M. smegmatis* NDH-2 and HQNO (*orange*) in *C. thermarum* NDH-2. Distances to the corresponding FAD co-factor are indicated. **B)** Comparison of *C. thermarum* NDH-2 residues that bind HQNO (*cyan*) and the corresponding residues in *M. smegmatis* NDH-2 (*purple*).

**Table S1:** Cryo-EM and model building statistics.

| | **NDH-2** (EMD-48544, PDB 9MQY) | **Compound 1-bound NDH-2** (EMD-48546, PDB 9MQZ) |
|---|---|---|
| **Data collection and processing** | | |
| Magnification | 120,000 | 165,000 |
| Voltage (kV) | 300 | 300 |
| Energy filter slit width (eV) | N/A | 10 |
| Electron exposure ($e^-/Å^2$) | 70 | 70 |
| Defocus range (µm) | 1.1 – 2.0 | 0.8 – 2.0 |
| Raw pixel size (Å) | 0.64 | 0.73 |
| Symmetry imposed | C2 | C2 |
| Initial particle images (no.) | 3,118,052 | 1,125,984 |
| Final particle images (no.) | 59,398 | 40,878 |
| Map resolution (Å) | 3.0 | 3.0 |
| FSC threshold | 0.143 | 0.143 |
| Map resolution range (Å) | 2.9 – 4.0 | 2.9 – 5.0 |
| **Model building and refinement** | | |
| Initial model used | AlphaFold-predicted model (AlphaFoldDB: AF-A0QYD6-F1-v4) | Inhibitor-free NDH-2 model |
| Model resolution (Å) | 3.1 | 3.4 |
| FSC threshold | 0.5 | 0.5 |
| Map sharpening *B* factor (Å) | –107.8 | –108.3 |
| **Model composition** | | |
| Nonhydrogen atoms | 6,287 | 6,157 |
| Protein residues | 894 | 886 |
| Ligands | FAD: 2 | FAD: 2, A1BNR: 2 |
| **R.m.s. deviations** | | |
| Bond lengths (Å) | 0.006 | 0.004 |
| Bond angles (°) | 0.697 | 0.810 |
| **Validation** | | |
| MolProbity score | 0.86 | 1.38 |
| Clashscore | 0.79 | 3.21 |
| Poor rotamers (%) | 0.67 | 0.71 |
| **Ramachandran plot** | | |
| Favoured (%) | 97.42 | 96.03 |
| Allowed (%) | 2.58 | 3.97 |
| Disallowed (%) | 0.00 | 0.00 |
